# Morphological determinants of glycosylation efficiency in Golgi cisternae

**DOI:** 10.1101/2025.11.14.688478

**Authors:** Christopher K. Revell, Nicola L. Stevenson, Martin Lowe, Oliver E. Jensen

## Abstract

The Golgi apparatus has an intricate spatial structure characterized by flattened membrane-bound compartments, known as cisternae. Cisternae house integral membrane enzymes that catalyse glycosylation, the addition of polymeric sugars to protein cargo, which is important for the trafficking and function of the products. The unusual and specific shape of Golgi cisternae is highly conserved across eukaryotic cells, suggesting significant influence in the correct functioning of the Golgi. Motivated by experimental evidence that disruption to Golgi morphology can lead to observable changes in secreted cargo mass distribution, we develop and analyse a mathematical model of polymerisation in a cisterna that combines chemical kinetics, spatial diffusion and adsorption and desorption between lumen and membrane. Exploiting the slender geometry, we derive a non-local nonlinear advection-diffusion equation that predicts secreted cargo mass distribution as a function of cisternal shape. The model predicts a maximum cisternal thickness for which successful glycosylation is possible, demonstrates the existence of an optimal thickness for most efficient glycosylation, and suggests how kinetic and geometric factors may combine to promote or disrupt polymer production.

**Author Summary:** The Golgi apparatus is a universal feature of eukaryotic cells, playing a critical role in post-translational modification of secreted proteins. Its importance is demonstrated by the variety of disorders caused by errors in its function. A major post-translational modification is glycosylation: the addition of sugar chains to protein cargo. The Golgi has a distinctive structure, being comprised of stacks of thin, hollow, flattened, cisternae. We develop a mathematical model of glycosylation, combining enzyme-mediated membrane-bound reactions with intra-cisternal diffusion, to quantify the effect of cisternal morphology on processing rate. The model identifies optimal and maximal cisternal thicknesses for glycosylation to proceed, in terms of biochemical parameters. The model offers a quantitative connection between the Golgi’s cisternal morphology and its function.

## Introduction

The Golgi apparatus is a critical organelle and a defining feature of eukaryotic cells [1–4]. It is typically comprised of between four and seven membranous compartments called cisternae, each of which have a distinctive flattened geometry reminiscent of a pita bread [5–7]. Each cisterna has a cylindrical diameter of roughly 1 µm and a depth of roughly 20-60 nm [8], indicating a spatial asymmetry of about 2 orders of magnitude that generates a high surface-area-to-volume ratio. This ratio is often further enhanced by the presence of fenestrations. The cisternae within the Golgi apparatus are categorised into three “maturation” states (*cis, medial*, and *trans*), each characterised by their enzymatic and biochemical composition. The *cis* cisternae are the entry point for cargo; *medial* cisternae are the intermediate maturation state; and the *trans* cisternae are the final compartments preceding cargo sorting and Golgi exit in the *trans*-Golgi network. Among competing theories of the cisternal maturation mechanism, one posits that stable Golgi cisternae exist with a given maturation state, and cargo is exchanged between them via transport vesicles [9], whereas another proposes that Golgi cisternae mature dynamically from one state to the next, driven by the recycling of Golgi resident proteins, whilst retaining their cargo and moving steadily through the stack, as though on a conveyor belt [10, 11]. Consensus seems to mostly favour the latter, but this does not affect our primary aim, which is to understand how cisternal shape influences Golgi function.

In humans, approximately 30% of all proteins pass through the Golgi. Any errors in the modification and trafficking of these products can lead to a wide variety of disorders [12–15], highlighting the importance of the Golgi apparatus for cellular function. Genes encoding proteins destined for the secretory pathway are transcribed in the nucleus to produce mRNA molecules bearing signal sequences that direct them to specific domains on the endoplasmic reticulum (ER). Here they form a template for the synthesis of protein chains and are co-translationally imported into the ER. After folding and post-translational modification, for example by disulphide bond formation or addition of N-linked glycans, proteins exit the ER and are transported to the Golgi apparatus where they are further modified and sorted for transport to their next cellular destination.

In most organisms, Golgi cisternae are stacked in close apposition with a *cis–trans* polarity established by directional transport. This stacking, whilst highly conserved, is not essential to survival as multiple independent events of unstacking can be observed in evolution, such as in the yeast *Saccharomyces cerevisiae* [16]. In many higher organisms, cisternae are also laterally connected to form a Golgi ‘ribbon’ architecture, a feature that evolved as a new functionalisation in the last common ancestor of cnidarians and bilaterians [17]. In contrast to stacking and lateral tethering, the overall geometry of a flattened cisterna has been universally conserved since the last eukaryotic common ancestor [3, 17]. It is therefore an ancient feature, implying a fundamental role in Golgi function. Despite this, very little is known regarding its functional relevance. This is due, in part, to experimental limitations, since the methods currently available to perturb cisternal flattening inevitably have pleiotropic impacts on Golgi biology. Genetic perturbation of the Golgi organising proteins GOLPH3 [18] or GMAP210 [19–23], or of the Golgi anion exchanger AE2 [24], will induce cisternal dilation, as will drug-induced disruption to actin polymerisation [25]. However, in each case other aspects of Golgi organisation, trafficking and chemistry are affected. Alternative approaches are therefore required to isolate this feature.

One of the major post-translational modifications orchestrated by the Golgi is the sequential addition of carbohydrate residues to cargo proteins to form glycan chains [26]. This process, called glycosylation [27], is mediated by enzymes anchored into the membrane of the Golgi cisternae via transmembrane protein domains, with the catalytic domain protruding into the lumen [28]. Unlike peptides, the synthesis of glycan structures is not templated and must be regulated by controlling conditions within the secretory pathway. This is achieved by controlling the cohort of enzymes expressed and their relative abundance, the density of sugar transporters and availability of monosaccharide substrates, the distribution of enzymes across the cisternal stack, the transport dynamics of cargo, and the spatial relationships between enzymes and substrates. As a result of these competing factors, the glycan profiles produced by any given cell population are generally heterogeneous. Existing studies are slowly beginning to unpick the relative contribution of many of these control points to glycosylation outcomes. However the role of Golgi morphology has remained particularly enigmatic.

In this study we use mathematical and computational modelling to test our prediction that cisternal shape is conserved to promote efficient glycosylation. The membrane-bound nature of the glycosylation reaction suggests that the Golgi apparatus would perform most efficiently with a high surface area to volume ratio, an intrinsic feature of the flattened morphology adopted by the cisternae. Other organelles with membrane-based functions, such as mitochondria, also have a high surface area to volume ratio, however this is achieved through alternative adaptations such as the folding of membranes into cristae.

Previous theoretical studies have examined a number of organisational features of the Golgi apparatus. For example, reaction networks expressed as stochastic and ordinary differential equations have been used to model the distribution of different branching oligosaccharide structures formed by glycosylation, and how this is influenced by the distribution of enzymes within the Golgi compartments [15, 29–32]. Other work has used transport modelling or agent-based models to compare competing theories of Golgi maturation [11, 33, 34], reaction networks to describe the self-organisation and maturation of Golgi compartments by vesicular exchange [35–40], or discrete-element methods [41] and energy minimisation [42–45] to model self-organised membrane structures that recapitulate Golgi morphology. However, the link between Golgi morphology and effective post-translational modification remains (to our knowledge) under-explored.

Figure 1A shows how the Golgi in a trafficking mutant (GMAP210 knockout, as characterized in [20]) is dilated relative to wildtype. The extent of dilation over a population of cells is quantified in Figure 1B. Figure 1C shows the differences in molecular weight distribution of an example glycosylated cargo, biglycan (BGN), in these cells. Biglycan is a proteoglycan comprised of a core peptide of 40 kDa mass modified with two linear glycosaminoglycan (GAG) chains which increase the mass of BGN to more than 250 kDa when fully modified, as seen here in WT cells. Undermodification in KO cells leads to an increase in a 150 kDa intermediate. Motivated by the data in Fig. 1 (obtained using procedures outlined in [20] and in Supplementary Information S1 Text), we explore below the potential relationship between cisternal shape and the molecular-weight profile of a Golgi product, bearing in mind that shape will not be the only determinant of the product’s profile.

**Fig 1.**
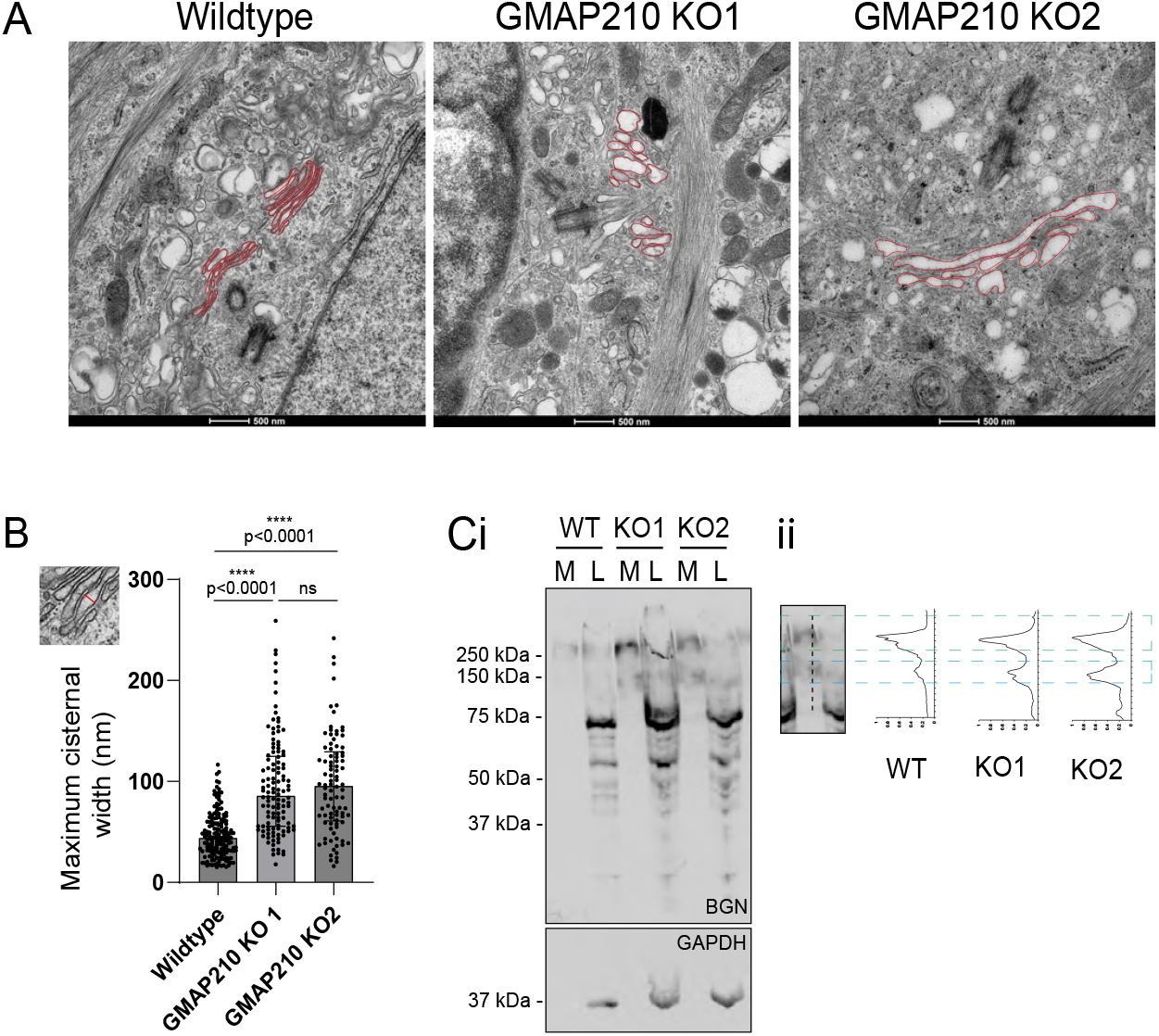
Experimental evidence suggests a link between cisternal dilation and biglycan undermodification in GMAP210 KO cells. **(A)** Transmission electron micrographs of wildtype (WT) and GMAP210 knockout (KO) RPE1 cells. Golgi membranes are outlined in red. **(B)** Quantification of the maximal depth of individual cisternae from images represented in A. The inset illustrates a measurement. Each dot represents one cisterna. In total 8-15 cells were measured across two independent experiments. All identifiable cisternae in each cell were measured. Data were analysed with a D’Agostino & Pearson test for normality (failed) and *p*-values were calculated using a Kruskall Wallis with Dunn’s multiple comparisons test. **(C)i** Western blot of media (M) and lysates (L) collected from cultures of wildtype and GMAP210 KO RPE1 cells stably expressing BGN-SBP-mScarlet-i. Samples were probed with antibodies targeting BGN and GAPDH as a loading control. Two KO clones are presented. **(C)ii** Line scan profiles of RFP fluorescence intensity through the media lanes of the blot represented in Ci. Green and blue dashed boxes show alignment of intensity peaks with bands on the blot in an example lane and line scan.

We model glycosylation within a cisterna as an enzyme-mediated polymerisation reaction (Fig. 2 and Eq. (S1) in Supplementary Information S2 Text). For simplicity, we consider primarily the assembly of linear polysaccharides, best represented biologically by the glycosaminoglycans and as represented in Fig. 1. Glycosaminoglygan synthesis initiates with the assembly of a core tetrasaccharide on a serine residue of a cargo peptide. This chain is then extended through the addition of repeating disaccharide units until the cargo exits the Golgi, with no concurrent polymer trimming [46]. Due to the repetitive nature of this structure, the overall chain length for each polymer is determined by numerous competing factors, such as transport time and enzyme concentration, resulting in the production of a heterogeneous mix of products. The molecular weight distribution of glycosylated output can therefore provide a sensitive read-out of glycosylation efficiency and its relationship with Golgi conditions including, as relevant here, changes to Golgi structure [13]. The polymer cargo and its monomeric substrate are present in the lumen of the cisterna but react only after adsorbing to a nearby cisternal wall, where a membrane-bound enzyme catalyses addition of a monomer to the polymer. The product of the reaction can desorb back into the lumen and the reaction can repeat. The polymer and substrate move within the lumen by molecular diffusion in a crowded environment. After some time we suppose that the contents of the lumen are released. In the framework illustrated in Fig. 2, the resulting polymer distribution can be expected to depend on four surface reaction rates, four adsorption/desorption rates of polymer and substrate, and geometric factors related to the size and shape of the cisterna. We wish to establish in particular how geometric factors may promote or hinder the glycosylation process.

**Fig 2.**
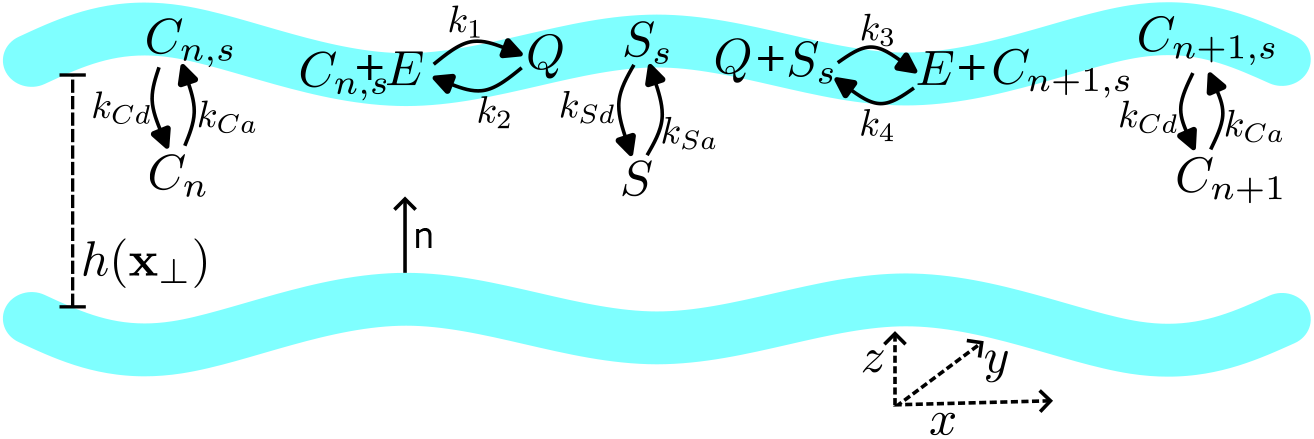
Schematic diagram of the mathematical model. Two cisternal membranes (blue) separate a lumen of thickness *h*(*x, y*); the cisterna lies in the (*x, y*)-plane and has inward normal n. Substrate molecules *S* and cargo molecules *C*_1_, *C*_2_, …, *C*_*N*_ diffuse freely within the lumen and adsorb reversibly onto the membranes to form membrane-bound substrate *S*_*s*_ and cargo *C*_*n,s*_. On each membrane, *C*_*n*_ binds reversibly to membrane-bound enzyme *E* to form membrane-bound complex *Q*, which then reacts with *S*_*s*_ to form *C*_*n*+1,*s*_, releasing the enzyme *E*. The cargo *C*_*n*+1,*s*_ is then free to reversibly desorb from the membrane back into the lumen. This system is characterised by 8 separate reaction rates, as labelled in the diagram.

## Materials and Methods

### The mathematical model

Full details of the mathematical model are presented in Supplementary Information S2 Text. Here we summarise the main assumptions and approximations and introduce the primary variables and parameters. Below, stars denote dimensional quantities.

The model describes adsorption of polymer cargo with luminal concentration 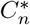 (where *n* = 1, 2, … describes the polymer length) onto cisternal membranes, where an enzyme-mediated polymerisation step takes place. This is followed by desorption of polymer with length *n* + 1 into the cisternal lumen, following the reaction scheme illustrated in Fig. 2 and given in Eq. (S1). The model combines the polymerisation reaction, modelled using mass-action kinetics, with transport by molecular diffusion in the cisternal lumen. The following set of approximations are exploited that permit substantial simplification of the model, facilitating its analysis.

1. The flat and slender geometry of the cisterna, which remains fixed, enables the bulk concentration 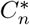 to be represented by a depth-averaged concentration.
2. Adsorption and desorption are assumed to be sufficiently rapid for 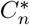 always to be in equilibrium with a surface concentration 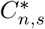. The affinity of cargo for the surface is captured by a lengthscale *h*_*C*_ ≡ 2*k*_*Ca*_*/k*_*Cd*_, a ratio of adsorption rate *k*_*Ca*_ to desorption rate *k*_*Cd*_. The factor of 2 is included because two surfaces bound the lumen (Fig. 2).
3. The (disaccharide) substrate, with luminal concentration *S*^∗^, is assumed to be sufficiently abundant for polymerisation not to change *S*^∗^ appreciably; the substrate is also assumed to adsorb rapidly to the membrane with an affinity captured by a lengthscale *h*_*S*_ ≡ 2*k*_*Sa*_*/k*_*Sd*_, a ratio of adsorption rate to desorption rate of substrate.
4. The surface concentration of membrane-bound enzyme, *E*^∗^, is assumed to be low enough to be rate-limiting.
5. By assuming that a large number of polymerisation steps take place, and following assumption 2 above, the polymer distributions 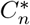 and 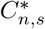 (with 1 ≤ *n* ≤ *N* for some *N* ≫ 1) can be represented by a dimensionless density 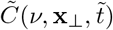 that depends on position **x**_⊥_ ≡ (*x, y*) in the plane of the cisterna, a dimensionless time 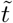 and a continuous variable *ν*, where 0 ≤ *ν* ≤ 1, that interpolates *n/N*.

Expressing the model in dimensionless form, and using thin-layer asymptotics [47] under assumptions 1–2 above, we reduce the diffusion equation in three space dimensions, describing luminal transport of polymer and substrate, to a spatially two-dimensional (2D) diffusion equation that is coupled to a set of ordinary differential equations that describe surface reactions using mass-action kinetics (Eq. (S16) in Supplementary Information S2 Text). Then, exploiting assumptions 3–5 above, the multiple ordinary differential equations of mass action kinetics can be combined with the bulk transport equation to recover the following partial differential equation, with polymerisation state represented as a continuous variable *ν*, coupled to an integral:

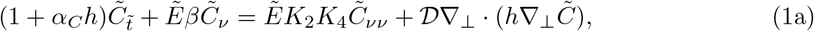

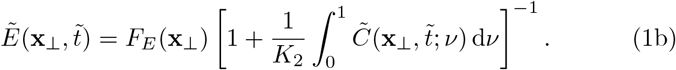

This is Eq. (S39) in Supplementary Information S2 Text. Subscripts *ν* and 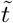 denote partial derivatives. Eq. (1a) is a non-linear non-local advection-diffusion equation that describes evolution over time 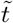 of the concentration field 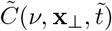 due to diffusion in 2D physical space **x**_⊥_ and advection-diffusion in polymerisation space *ν*.

The first term of Eq. (1a) represents the rate of change of the combined surface and luminal concentrations; surface reactions are captured by *ν* derivatives and diffusion in the lumen is represented by the final term in Eq. (1a). Since enzyme at any location binds with molecules of any molecular weight, its available (dimensionless) concentration 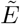 is given by the integral in Eq. (1b). The model accommodates heterogeneities in cisternal depth *h*(**x**_⊥_) (via the capacitive prefactor of the time derivative 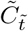 and via non-uniform spatial diffusion) and in enzyme distribution (through *F*_*E*_(**x**_⊥_) in Eq. (1b)), for some given functions *h* and *F*_*E*_; these satisfy the constraints ∫ *h*(**x**_⊥_) d**x**_⊥_ = *π*, ∫ *F*_*E*_(**x**_⊥_) d**x**_⊥_ = *π*, when integrated over the cisterna. Parameters *α*_*C*_, *K*_2_, *K*_4_ in Eq. (1) are defined in Table S2 of Supplementary Information S2 Text; parameters *β* and *D*, which combine biochemical and geometric quantities, are defined in Eq. (S34) and Eq. (S40) respectively. *α*_*C*_ compares the mean cisternal thickness to the adsorption lengthscale *h*_*C*_; the term proportional to *α*_*C*_*h* in Eq. (1a) represents polymer residing in the cisternal lumen, and is the primary means by which cisternal geometry influences polymerisation rate. *β >* 0 measures the strength of the forward polymerisation reaction. The product *K*_2_*K*_4_ measures reverse reaction rates (illustrated by *k*_2_ and *k*_4_ in Fig. 2) that contribute to dispersion in molecular weight as polymerization proceeds (arising because 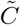 describes a population of polymers that are effectively undergoing a biased random walk in *ν*-space). *D* measures spatial diffusion in the plane of the cisterna.

Starting with a concentration field derived from a Gaussian distribution with small variance, representing monomeric cargo at 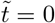, we simulate Eq. (1) until some release time 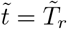. Integration of 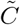 over the spatial domain then yields 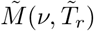, a distribution of polymer with respect to scaled molecular weight *ν* at the release time. Treated as a function of *ν*,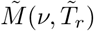 serves as a proxy for the result of a Western blot experiment (as in Fig. 1Cii). We treat the luminal mass of polymer having molecular weight satisfying 0 *< ϕ < ν* ≤ 1, for some *ϕ*, as a measure of the functional output of the cisterna. The dimensional mass of luminal polymer having chain length *n* in the range *ϕN* ≤ *n* ≤ *N* at time 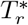 (Eq. (S4) in Supplementary Information S2 Text) is

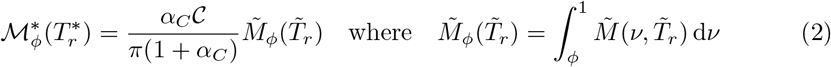

(Eq. S44 in Supplementary Information S2 Text). Here *C* is the initial number of cargo monomers in the cisterna. To quantify cisternal efficiency, we evaluate the time 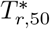 taken to produce 50% of the maximum possible functional output of the reaction, were it allowed to run indefinitely, and define the production rate 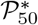 as this functional polymer mass in the cisternal lumen divided by 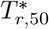 (Eq. (S5)). The primary aim of the model is to explore how different cisternal shapes *h*(*x, y*) in Eq. (1a), in combination with kinetic parameters, influence the production rate 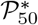.

The input parameters of the model include 8 rate constants (Fig. 2; Table S2 of Supplementary Information S2 Text) and three chemical masses per cisterna (of monomer *C*, substrate *S* and enzyme ℰ; Table S4). Table S7 of Supplementary Information S2 Text reports parameter values used in computations. We necessarily make assumptions about the relative sizes of some of these quantities, in order to reduce the model to a tractable and interpretable form. We recognize that direct evidence for many of these assumptions is not currently available and that model predictions must therefore be interpreted cautiously. Values of input parameters in Table S7 are given in terms of a nominal reference length L = 100 nm (comparable to a cisternal width, as in Fig. 1B), a time T = 10^−5^ s (representative of a desorption rate) and a molecular number M = 10^3^ (representative of a scarce number of enzyme molecules in a cisterna), that we choose in order to give reasonable outcomes. Time 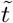 is scaled as shown in Table S4, so that 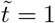 corresponds to 2600 s (43 min) for the illustrative parameters chosen in Table S7; the spatial coordinate **x**_⊥_ is scaled as shown in Table S3; dimensional luminal and surface concentrations are recovered from 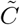 via

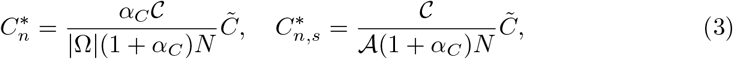

where |−| is the luminal volume and *A* ≡ 2|Ω_⊥_| is the total cisternal membrane area. The simplifications described above substantially reduce the number of independent dimensionless parameters (summarized in the right-hand columns of Table S2 and S4), that are implemented in Eq. (1). In summary, by retaining processes that are likely to be dominant, the model reveals specific dimensionless combinations of parameters that determine outcomes, and provides plausible quantification of the contributions of different physical and chemical processes to glycosylation in a cisterna.

### Computational method

A full explanation of computational techniques used to solve Eq. (1) is provided in Section S3 of Supplementary Information S2 Text, along with code needed to reproduce results in this paper. All numerical results were generated using code written in the Julia programming language [48], leveraging the `DifferentialEquations.jl` library [49], a strong stability-preserving Runge–Kutta solver [50], the `Makie.jl` plotting library [51], and an incidence matrix approach to differential operators [52]. Unless otherwise specified, parameters as laid out in Table S7 were used for all results.

## Results

While there is uncertainty in the values of many of the input parameters, the model offers predictions of production rate that reveal parameter dependencies and which give insight into factors that promote or hinder functional polymer production.

### Depth-dependence of production rate

We first illustrate polymerisation in a cisterna of uniform thickness *h*_0_ (corresponding to a dimensional thickness *h*_0_L), having spatially uniform concentration distributions. (Analytic model predictions for this case are described in Section S4 of Supplementary Information S2 Text.) Fig. 3A shows how the polymer concentration distribution, initially monomeric (so that 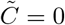 at 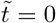 away from *ν* = 0), takes on a Gaussian profile that propagates towards *ν* = 1 as polymerization proceeds. The mass of functional polymer 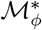 (see Eq. (2)) is determined from its dimensionless equivalent 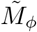), which rises from zero to *π* (Fig. 3B) as the 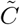 distribution sweeps past *ν* = *ϕ*, allowing us to identify the half-rise time 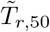. (The time 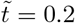 translates to approximately 9 minutes, using the parameters in Table S7.) 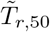 is converted to a dimensional 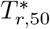 (via Eq. (S48)) and used to estimate production rate 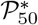 as shown in Fig. 3C. The production rate magnitude in this example, 0.89 *×* 10^−4^ M*/*T, which is equivalent to approximately 9000 molecules/s for our choice of M and T (see the black cross in Fig. 3C), arises from the specific choice of parameter values in Table S7. 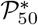 can be measured relative to the factor *k*_1_*C*ℰ*/*(*AN*), which reflects the strength of the initial forward reaction and takes the value 10^−2^M/T = 10^6^ molecules/s in this example; here the enzyme concentration is estimated via ℰ*/*(*A*. The factor *N* reflects the requirement for this reaction to repeat many times to form a functional polymer. *k*_1_*C*ℰ*/*(*AN*) overestimates the production rate because the second forward reaction rate *k*_3_ (Fig. 2) is small (*k*_3_ = *k*_1_*/*40) in this example (Table S7).

**Fig 3.**
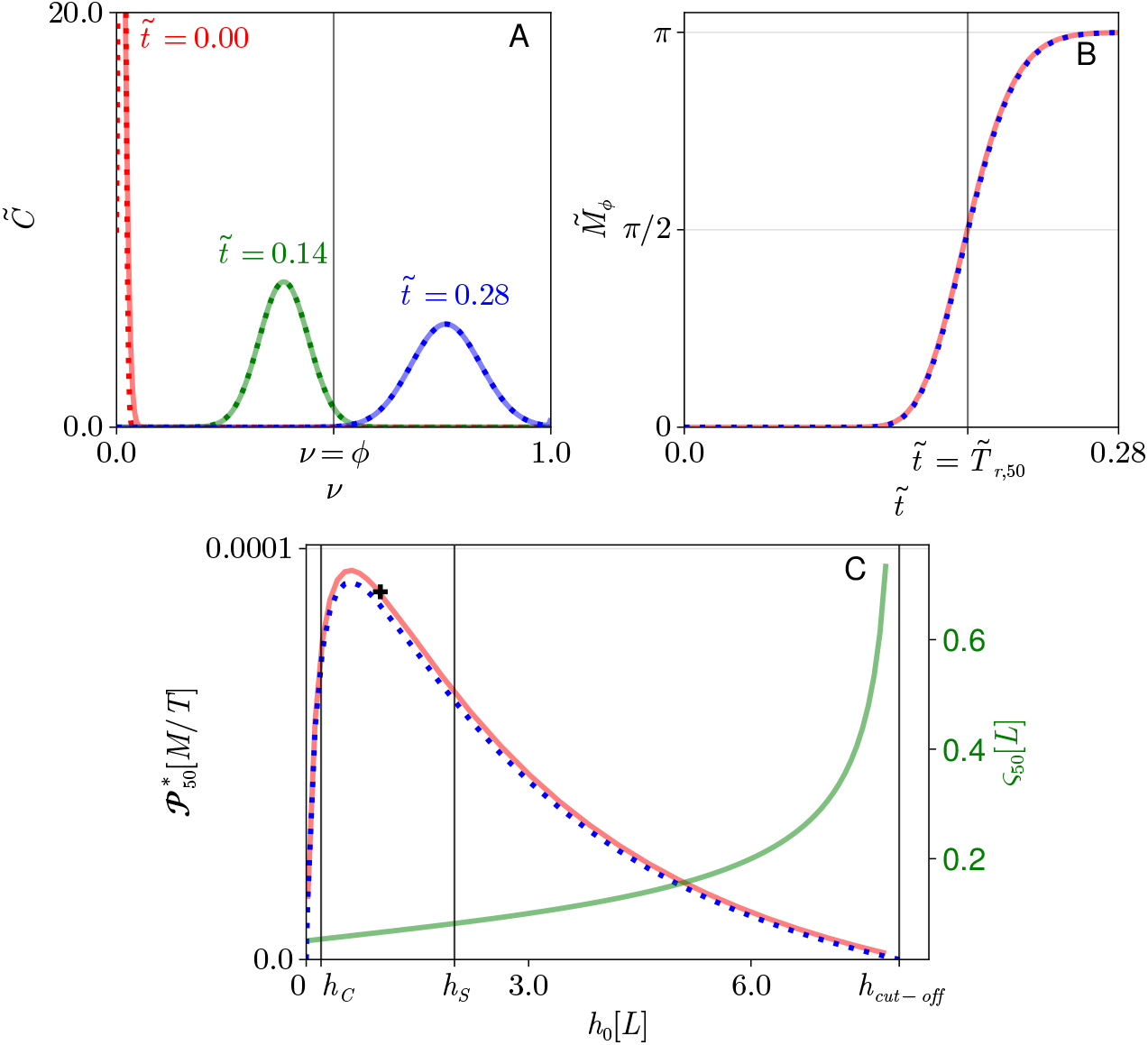
Polymerization in a cisterna of uniform depth. **(A)** Solutions of Eq. (1) assuming uniform cisternal thickness and enzyme distribution, showing dimensionless polymer concentration distribution 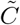 versus polymerisation length *ν* = *n/N* at three times, indicated by red, green and blue lines. Numerical solutions (solid lines) are compared to the analytic solution (Eq. (S53); dashed lines) with 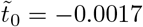 and *ν*_0_ = 0.0080. The vertical line shows the threshold *ν* = *ϕ* above which polymer is treated as functional. **(B)** Functional polymer production 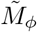 (see Eq. (2)) against time 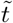, comparing numerical result (solid line) against analytic result (Eq. (S54); dashed line). The time to produce half of the maximum possible production, 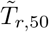, is shown. **(C)** Left axis: Dimensional production rate 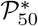 of functional cargo versus cisternal depth *h*_0_, comparing numerical result (Eq. (S50); solid line) to analytic result (Eq. (S58); dashed line). *h*_*C*_ and *h*_*S*_ are adsorption depths of cargo and substrate. *h*_cut−off_ is given in (5). Right axis: standard deviation *ς*_50_ of 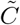 at 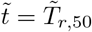 from (S68, S69) in Supplementary Information S2 Text. Parameters are given in Table S7; units of 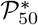 and *h*_0_ are given in terms of reference quantities M, L and T. Peak production rates are 0.947 × 10^−4^ at *h*_0_ = 0.62 in numerics, and 0.917 × 10^−4^ at *h*_0_ = 0.72 in analytic expression. The black cross shows the production rate 0.895 × 10^−4^ at *h*_0_ = 1, corresponding to panels A and B.

A flat cisterna allows the governing partial differential equation to have an analytic solution (Eq. (S53) in Supplementary Information S2 Text), which agrees closely with simulations (Fig. 3A, B) after introducing two fitting parameters linked to details of the initial conditions used in computations. The values of these fitting parameters are very small, so disregarding these corrections, the analysis provides a simple expression for production rate of the form

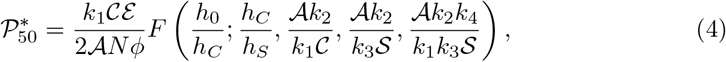

where the function *F* is a ratio of polynomials in *h*_0_*/h*_*C*_ (Eq. (S58)). The four dimensionless groups that appear as *h*_0_-independent parameters in *F* measure the relative adsorption strength of substrate and polymer (*h*_*C*_*/h*_*S*_), and their abundances when adsorbed over the total cisternal membrane area *A*. *k*_1_ and *k*_3_ (*k*_2_ and *k*_4_) are forward (reverse) reaction rates (Fig. 2); the reaction requires *Ak*_2_*k*_4_*/k*_1_*k*_3_*S* to be sufficiently small in order to proceed. Equation (4) captures the major qualitative features of simulations (Fig. 3A,B) and reveals direct dependencies of 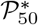 on input parameters. We focus here on its variation with cisternal depth, *h*_0_ (Fig. 3C).

The production rate initially rises from zero for increasing cisternal depth *h*_0_ (Fig. 3C). This is because functional polymer is assumed to reside in the lumen, in equilibrium with that adsorbed onto membranes, and a very thin cisterna accommodates proportionally less functional polymer. The production rate reaches a maximum value as depth increases before falling to zero at a cut-off depth defined by

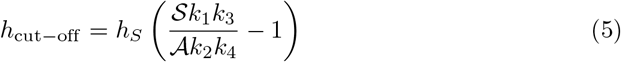

(Eq. (S36) in Supplementary Information S2 Text). For *h*_cut−off_ to be positive requires the substrate to be sufficiently concentrated on the cisternal membrane, requiring *S/A* to exceed the ratio *k*_2_*k*_4_*/k*_1_*k*_3_ between the forward and reverse reaction rates (Fig. 2). The cut-off thickness is proportional to the substrate adsorption depth *h*_*S*_.

Equation (4) can be used to identify the cisternal depth that maximises the production rate. It is helpful to evaluate this optimum in limiting cases. For example, when *h*_*C*_ ≪ *h*_*S*_ (as in Fig. 3), implying that the polymer adsorbs to the membrane less readily than the substrate, the optimal cisternal thickness can be written

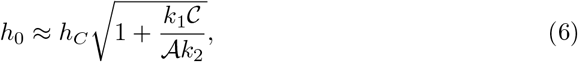

leading to maximum production rate given by Eq. (S62) in Supplementary Information S2 Text. Eq. (6) predicts an optimum depth of 0.92 L, consistent (to this level of approximation) with the peaks lying at 0.62 L (solid curve) and 0.72 L (dashed curve) in Fig. 3C. The optimal cisternal thickness in this case is regulated by the adsorption properties of the polymer (*h*_*C*_), mediated by the abundance of polymer. If in addition the cargo is sufficiently abundant, so that the *C* term dominates in Eq. (6), the associated maximum production rate (Eq. S63) is limited by polymer and enzyme abundance.

Figure 3C also shows how the standard deviation *ς*_50_ of the 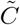 density at 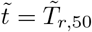 (Eq. (S68, S69)) grows with cisternal depth, becoming unbounded as *h*_0_ approaches *h*_cut−off_. In addition to slowing the reaction, dilution of substrate provides more opportunity for reversals of the polymerization process that disperse the distribution. Thinner cisternae therefore promote a purer product with a more narrowly defined molecular-weight distribution.

In summary, the model (as illustrated by Fig. 3) shows how, for a cisterna of uniform depth *h*_0_, the functional polymer production rate is regulated by cisternal depth: the depth must be below *h*_cut−off_ for the substrate to be sufficiently concentrated for the forward reaction to proceed; and the affinities of solute and substrate for the membrane (as expressed by the thicknesses *h*_*C*_ and *h*_*S*_) determine the thickness at which polymer production proceeds most rapidly. Unfortunately, given uncertainties in model parameters, it is not possible to make definitive predictions of optimal cisternal depth for specific cargoes; however, expressions such as Eq. (6) (or more general expressions that can be derived from the function *F*) reveal the factors that regulate optimal width in terms of the dominant biochemical and transport processes.

### Dispersion driven by depth and enzyme heterogeneity

There is intrinsic dispersion in the polymerisation process, regulated by biochemical rate constants and evident in Fig. 3A. This is captured in the model by the term representing diffusion with respect to *ν* in Eq. (1). However, dispersion can also arise from spatial heterogeneities, either in cisternal shape or in enzyme distribution, as illustrated in Fig. 4.

**Fig 4.**
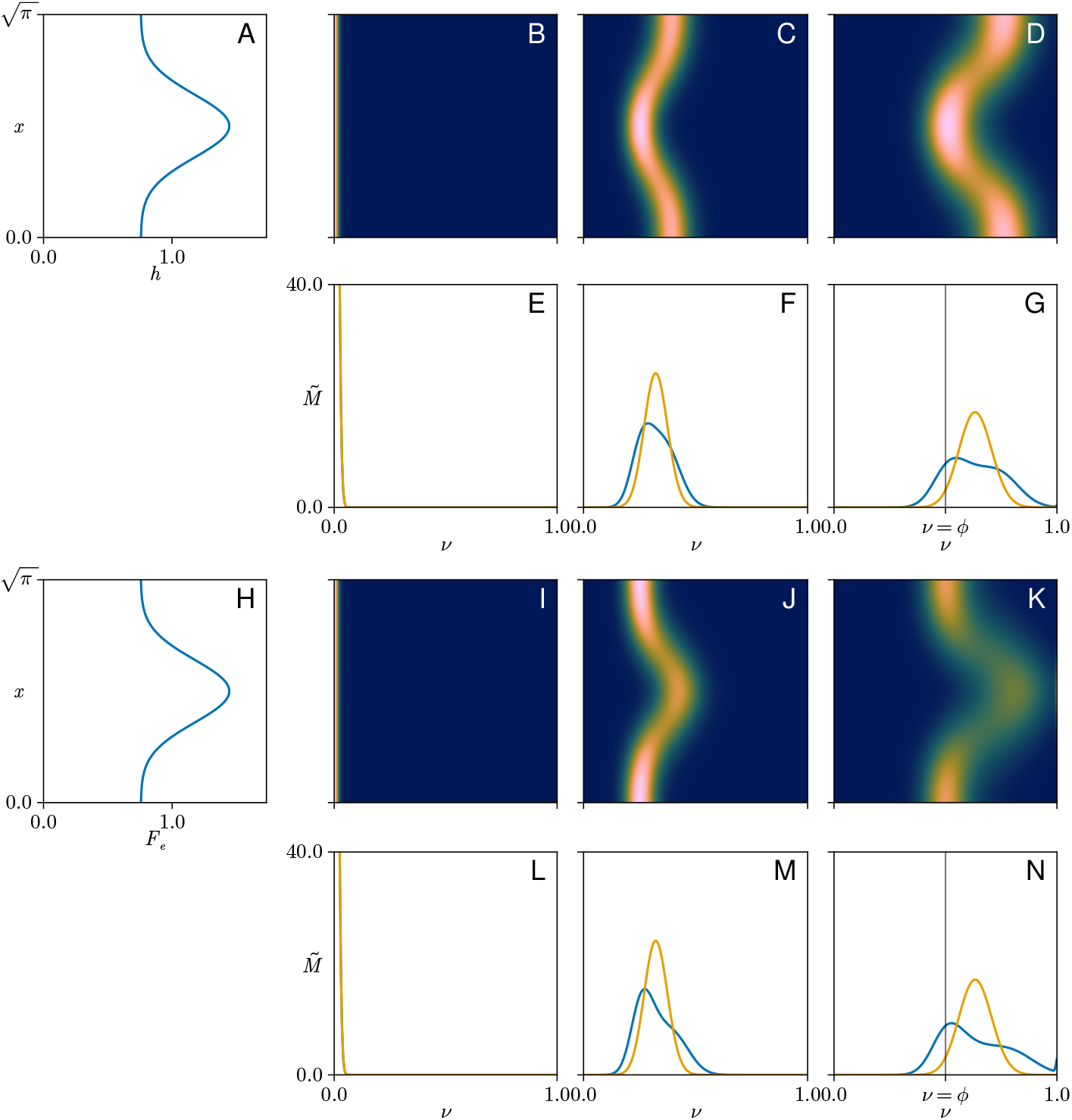
Non-uniformities in one spatial dimension. **(A)** Dimensionless cisternal thickness *h*(*x*) against *x* showing a Gaussian thickening. **(B**,**C**,**D)** Heatmaps of concentration 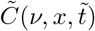 in real (vertical) and polymerisation (horizontal) space at initial, intermediate, and final states (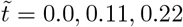) associated with the thickness profile in A. **(E**,**F**,**G)** Blue curves: spatial integral of concentration, 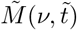, corresponding to B-D, versus polymerisation length *ν*. Orange curves show 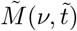 when *h* = 1 and *F*_*E*_ = 1 uniformly. **(H)** Gaussian enzyme distribution *F*_*E*_(*x*) against *x*. **(I**,**J**,**K)** Heatmaps of concentration 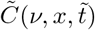 in real and polymerisation space at initial, intermediate, and final states (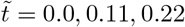) given the enzyme distribution in (H) and assuming spatially uniform *h*(*x*) = 1. **(L**,**M**,**N)** Blue curves: spatially integrated concentration distributions, 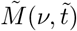, corresponding to I-K. Orange curves show 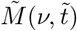 when *h* = 1 and *F*_*E*_ = 1 uniformly. Parameters found in Table S7. Vertical lines in G, N show the threshold *ν* = *ϕ* above which polymer is regarded as functional.

To illustrate the role of spatial heterogeneity, we consider a cisterna with a localised region of increased depth, having a profile as shown in Fig. 4A. (Here, for illustrative purposes, we consider variation with respect to a single spatial dimension and symmetry in the second dimension; this example is equivalent to a cisterna having a straight ridge of increased thickness.) When Eq. (1) is solved for this depth profile and a spatially uniform initial monomer distribution (Fig. 4B), we see significant spatial variation in polymerisation of cargo (Fig. 4C,D). Since surface-bound polymer is in equilibrium with luminal polymer, the surface reaction leads to changes in the combined population integrated across the cisterna with mass proportional to 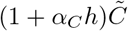 per unit area (see the first term in Eq. (1a)). It follows that polymerisation proceeds quicker where the cisterna is thinner; equivalently, thicker regions hold more polymer, requiring longer for this polymer to be processed). Integration over space (to obtain 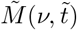 shows how the polymer mass distribution may become bimodal (Fig. 4E-G). The model therefore illustrates how spatial heterogeneity promotes variation in the secreted polymer mass distribution. Furthermore, spatial heterogeneity reduces the production rate 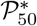 slightly in this example, from 0.895 *×* 10^−4^ (when *h*_0_ = 1, as in Fig. 3A,B) to 0.872 *×* 10^−4^ in Fig. 4A-G.

A further source of dispersion arises from heterogeneity of enzyme distribution. For example, enzymes have been shown to preferentially localise towards the cisternal interior [53]. Suppose, therefore, in a cisterna of uniform depth, that there is a region where enzyme is locally enriched, as illustrated in Fig. 4H. Since enzyme availability is rate-limiting in the present model, polymerisation proceeds more rapidly in regions of higher enzyme concentration (Fig. 4I-K). Spatial integration leads once again to bimodal polymer mass distributions (Fig. 4L-N), again with a marginal reduction in production rate compared to the uniform case (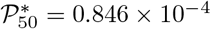, compared to 0.895 *×* 10^−4^ when *h*_0_ = 1 and *F*_*E*_ = 1).

### Random domains

The examples in Fig. 4 have variation in one spatial dimension but are symmetric in the second spatial dimension, and as such are illustrative of mechanisms but lack the spatial disorder evident in real cisternae. To accommodate this, we simulated polymerisation with depth variation derived from a 2D Gaussian random field [54, 55], as shown in Fig. 5A. The spatial domain is chosen for convenience to have a square footprint. Numerical integration of Eq. (1) gives 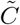 as a function of *ν*, **x**_⊥_ and 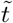. Visualisation of a scalar field over four dimensions is difficult, so we present projections of the concentration distribution 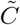 in Fig. 5B. Again, polymerization proceeds more quickly where the cisterna is thinner. We can further evaluate 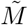 and compare to a uniform-depth case with the same parameters, as in Fig. 5C. This again demonstrates how spatial variation in cisternal depth leads to a broader distribution of cargo mass.

**Fig 5.**
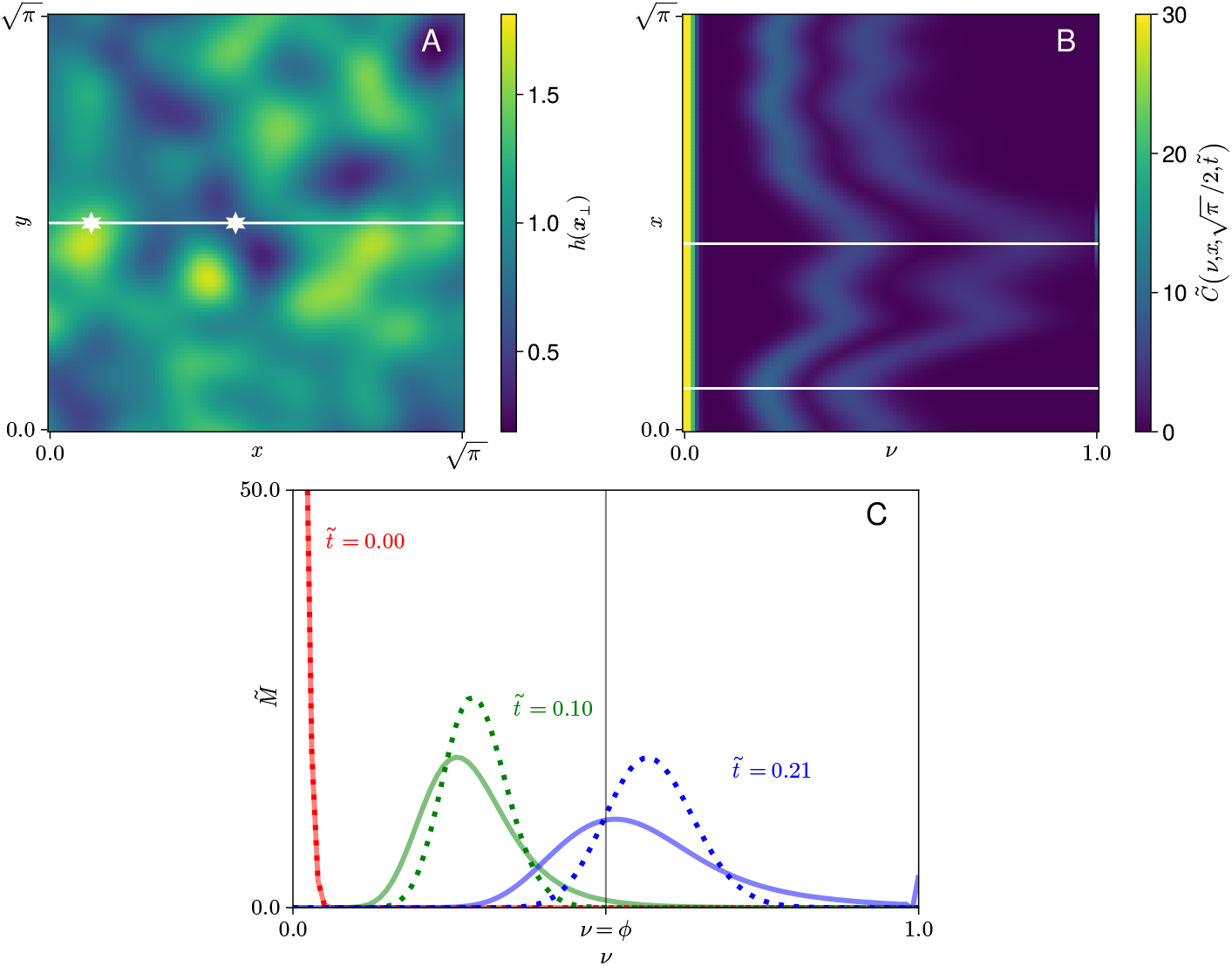
Nonuniformities in two spatial dimensions. **(A)** Cisternal thickness profile from Gaussian random field with mean 1, standard deviation 0.3, and correlation length The white horizontal line shows the slice at *y* = *π/*2 used in panel B; points of minimum and maximum thickness on the line are shown by white stars. **(B)** Heatmaps of 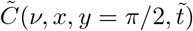 at times 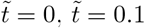 and 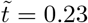 heatmaps are overlaid to demonstrate the evolution of the field. White lines show the locations of minimum and maximum *h*(*x, y* = *π/*2), as highlighted in panel B with white stars, corresponding respectively to regions of faster and slower polymerization. **(C)** Spatial integral 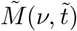 at the same three timepoints shown in panel B, with comparison to the uniform thickness case. The vertical line shows *ν* = *ϕ*, the threshold above which polymer is regarded as functional. Parameters found in Table S7.

## Discussion

The key conserved feature of Golgi morphology across all species and cell types is the structural flattening of cisternae into thin hollow discs. Such evolutionary conservation highlights this feature as pivotal to the function of the Golgi. The model presented in this study seeks to quantify the relationship between cisternal morphology and some of the biochemical processes underpinning the fundamental role of the Golgi in enzymatic protein modification. We have studied a canonical polymerisation process in some simple geometries in order to highlight dominant relationships.

Our strategy has been to reduce the model where possible to simple relationships, deriving explicit relationships that capture prominent features. The 17 input parameters of the full model were reduced to 11 dimensionless ratios by exploiting the small cisternal aspect ratio, and then to 6 dimensionless ratios by a set of additional biochemical assumptions. Despite uncertainties in the values of these quantities, we have been able to draw some noteworthy conclusions. For example, we have demonstrated that, subject to the model’s assumptions, an optimal cisternal depth is predicted that maximises the efficiency of useful glycosylation (Figure 3C); the optimal depth depends on biochemical parameters. (The Golgi processes multiple cargoes, so the optimum depth that we have evaluated for one species should be considered alongside the optima for others.) We have also shown that there exists a cut-off depth beyond which polymerisation is not predicted to proceed. Above the optimal depth, thickening of the cisterna leads to dilution of the available substrate, slowing polymerization, until the cut-off depth is reached at which polymerization ceases. These relationships are captured in the rational function *F* Eq. (4) (see Eq. (S58) in Supplementary Information S2 Text), which can be interrogated directly or which can be approximated in suitable limits. We illustrated in Eq. (6) the optimal thickness when the substrate adsorbs to the membrane more avidly than the polymer cargo, showing how the adsorption depth of the species that adsorbs more weakly to the membrane is a key determinant of optimal thickness. Equivalent expressions can be derived for other special cases, such as when the substrate adsorbs more strongly to the membrane than the cargo (Eq.(S66) in Supplementary Information S2 Text).

Reversibility of the individual steps in a polymerization reaction, such as those illustrated in Fig. 2, leads to dispersion in the molecular-weight profile of polymers sampled at a given time. This dispersion is captured by the *ν*-diffusion term in (1a). In a uniform cisterna, dispersion is predicted to increase dramatically with cisternal depth (Fig. 3C) as a result of substrate dilution. We have shown how dispersion is further enhanced in cases of spatial heterogeneity, at least under the assumptions of the model. Polymerisation proceeds more rapidly in thinner regions of the cisterna because, with substrate abundant and enzyme scarce relative to cargo, there is less cargo to process in a given time. Thus, when aggregated over a domain, the overall molecular weight profile broadens (Fig. 5) and can even become bimodal when the domain has distinct thick and thin regions (as illustrated in Fig. 4). Furthermore, we found that spatial heterogeneity reduced the production rate of functional polymer. Uniform cisternal thickness therefore promotes both the rate and the quality of glycosylated product, with purer product being achieved in thinner cisternae. Comparing the relatively uniform, thin morphology of wild-type Golgi with more heterogeneous and thicker cisternae of GMAP210 KO cells (Fig. 1A,B), the model may explain some of the changes in molecular weight distribution of glycosylated output from cells with aberrant Golgi (Figure 1C). We recognise however that the depth-dependent mechanism described here is only one of many potential causes of a widened molecular weight distribution. We showed for example that heterogeneous enzyme distribution can lead to similar outcomes.

The present model rests on numerous assumptions. This is a deliberate compromise, allowing us to capture key processes while maintaining a degree of tractability. Further stages of this work could implement changes to address some of these simplifications and assumptions. For example, we have adopted a simple model of O-glycosylation that ignores numerous complexities [56]. We show in Section S6 of Supplementary Information S2 Text how the model can be adapted for a peptide with two GAG chains such as biglycan, accommodating the new structure via a doubling of the coefficients of 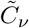 and 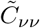 terms in Eq. (1a). Furthermore, we have ignored the possibility that adsorption and diffusion rates may depend on polymer size; it could be anticipated that longer polymer chains would be less mobile, and therefore less accessible to polymerisation, potentially placing an effective upper limit on polymer length. Numerous additional factors could be addressed, including stochastic fluctuations (arising from Brownian effects and small molecular numbers), the role of fenestrations, the off-target impacts of drug treatments affecting other cell biological processes such as Golgi transport, and alternative protein modifiications catalysed by membrane-anchored enzymes such as glycan-trimming, sulphation and phosphorylation. We acknowledge that, given the simplifications used here, the conclusions drawn from our model apply primarily to the chemistry taking part in the flattened central areas of the cisternae, and not the cisternal rims where cisternae are often more dilated. We believe this approach is valid in light of experimental studies demonstrating the spatial separation of Golgi transport and processing functions to the cisternal rims and flattened domains respectively [53]. Indeed, our findings provide a rationale supporting this model of domain functionalisation within the compartment.

In summary, we have presented a mathematical model, inspired by experimental data showing changes in the molecular weight distribution of glycosylated output from cells with changes in the thickness of their Golgi apparatus, that demonstrates an optimal thickness for glycosylation efficiency, and shows how heterogeneity in Golgi thickness leads to dispersion in, and hence potentially a reduction in precise control over, the molecular weight of glycosylated cargo.

## Supplementary information

**S1 Text. Supplementary text S1** describes procedures used to generate images in Fig. 1.

**S2 Text. Supplementary text S2** describes details of the mathematical model.

## Acknowledgements

This work was supported by the Biotechnology and Biological Sciences Research Council (BB/ T001984/1) and by Medical Research Council Award number UKRI1419.

For the purpose of open access, the authors have applied a Creative Commons Attribution (CCBY) licence to any Author Accepted Manuscript version arising.

Code used to generate these results can be found on GitHub [57].

## Author contributions

**Conceptualization**: all authors;

**Formal analysis**: CKR, OEJ, NS;

**Funding acquisition**: OEJ, ML;

**Investigation**: all authors;

**Methodology**: CKR, OEJ, NS;

**Project administration**: CKR, OEJ, NS;

**Resources**: NS;

**Software**: CKR;

**Supervision**: OEJ

**Visualization**: CKR

**Writing - original draft**: CKR, OEJ, NS;

**Writing - review and editing**: all authors.

