## Supplementary text S1 for "Morphological determinants of glycosylation efficiency in Golgi cisternae"

### Morphological determinants of gylcosylation efficiency in Golgi cisternae: Supplementary Information S1 Text

November 14, 2025

#### Cell culture

Telomerase immortalized human retinal pigment epithelial cells (hTERT-RPE1) were originally purchased from the American Type Culture Collection (ATCC). The generation and characterisation of the GMAP210 knockout lines is detailed in [1]. For culture, cells were grown in DMEM-F12-HAM (# 11320-033; Life Technologies) supplemented with 10% decomplemented fetal bovine serum (FBS, # A5256701; Gibco) at 37 °C/5% CO 2. Cells were passaged 1:10 every 3–4 days using 0.05% Trypsin-EDTA (Thermo-Fisher Scientific). Stable cell lines expressing BGN-SBP-mSc were also generated in [1]. In brief, lentivirus a carrying a pLVX-Neo-BGN-SBP-mSc plasmid (addgene # 234884) was made using the Lenti-X Packaging Single Shots kit (# 631275; Takara) according to kit instructions and used to transduce cells in the presence of 8 µg mL$^{-1}$ polybrene (# sc-134220; Santa Cruz Technology). Transfected cells were selected with 400 µg mL$^{-1}$ G418 geneticin disulphate.

#### Secretion assays

Cells were seeded in 6-well dishes and grown to confluence. The growth medium was then aspirated and replaced with 1 mL serum-free DMEM-F12-HAM to collect secreted proteins. After overnight incubation at 37 °C/5% $CO_2$, cells were put on ice and the media were collected, centrifuged at 2000g for 2 minutes at 4 °C to remove dead cells. The supernatants were collected and stored on ice to proceed with SDS-PAGE. Meanwhile the cell layer was washed with ice-cold PBS and lysed on ice for 15 minutes on a rocker in 100 µL RIPA buffer (50 mmol Tris-HCl, pH 7.5, 300 mmol NaCl, 2% Triton X-100, 1% deoxycholate, 0.1% SDS, 1 mmol EDTA) containing protease inhibitors (# 539137; Millipore). Lysates were scraped, collected up and centrifuged for 10 minutes at 13000g/4 °C. The supernatant was collected and stored on ice to proceed with SDS-PAGE.

#### SDS-PAGE and Western blotting

Media and lysates secretion assay samples were mixed with 4× Bolt LDS sample buffer (# B0007; Invitrogen) and boiled for 10 minutes at 70 °C. Samples were loaded on Bolt 4–12% Bis-Tris gels (Thermo Fisher Scientific) in MOPS running buffer (# P6010055; Thermo Fisher Scientific) and current applied for 40 minutes at 130 V. Proteins were transferred onto Amersham protran 0.2 µmol nitrocellulose membranes (# GE10600001; Merck) at 90 mA for 90 minutes. Membranes were blocked with 5% milk/TBST for one hour, then incubated with primary antibody overnight at 4 °C. Antibodies used were mouse anti-biglycan (abcam; ab188508; lot Gr2a5102.3) and anti-GAPDH (Proteintech; 60004-1-Ig; Lot 10028230). Membranes were washed with TBS/0.05% Tween (Sigma-Aldrich) and incubated with DyLightTM fluorophore-conjugated (Invitrogen) secondary antibodies in the dark for 2 hours prior to washing again. Blots were imaged using a Li-COR Odyssey system. Line profiles of fluorescent intensity were generated using the plot profile function of image J software.

##### Electron microscopy

Cells were grown to confluence in 35 mm dishes then fixed in 2.5% glutaraldehyde for 20 minutes, washed for 5 minutes in 0.1 mol cacodylate buffer and post-fixed in 1% OsO4 in 0.1 mol cacodylate buffer for 30 minutes. Cells were washed three times with water prior to staining with 3% uranyl acetate for 20 minutes. After another rinse with water, cells were dehydrated by sequential 10 minute incubations with 70%, 80%, 90%, 96%, 100% and 100% ethanol before embedding in Epon at 60 °C for 72 hours. Thin 70 nm serial sections were cut using a Leica UC6 ultramicrotome and a diamond knife. Sections were stained with 3% uranyl acetate then lead citrate, washing three times with water after each. Once dried, sections were imaged using an FEI Tecnai 12 120 kV BioTwin Spirit Transmission electron microscope. Cisternal depth was measured using imageJ software. Statistical tests were performed using GraphPad Prism software. For each condition 8-15 cells were imaged and measured across two independent experiments. To ascertain non-parametric distribution, samples were subjected to a D'Agostino & Pearson test for normality. As data were not normally distributed, p values were calculated using a Kruskall Wallis with Dunn's multiple comparisons test.
