## Supplementary text S2 for "Morphological determinants of glycosylation efficiency in Golgi cisternae"

### Morphological determinants of gylcosylation efficiency in Golgi cisternae: Supplementary Information S2 Text

November 14, 2025

Section S1 of this Supplement presents the full dimensional model describing glycosylation. A simplified form of the model is derived in Section S2. Section S3 explains the numerical scheme used to compute solutions of the simplified model. Results for a cisterna of uniform depth are given in Section S4. Parameter values are discussed in Section S5. Section S6 explains how the model can be adapted to described the evolution of two-chain polymers.

#### S1 The full model

Consider the polymerization reaction system for cargo $C_n^*$, enzyme $E^*$, complex $Q_n^*$, substrate (a saccharide) $S^*$, yielding product $C_{n+1}^*$:

$$S^* \underset{k_{Sd}}{\overset{k_{Sa}}{\rightleftharpoons}} S_s^*, \qquad C_n^* \underset{k_{Cd}}{\overset{k_{Ca}}{\rightleftharpoons}} C_{n,s}^*, \qquad n = 1, 2, \ldots, N, \tag{S1a}$$

$$C_{n,s}^* + E^* \underset{k_2}{\overset{k_1}{\rightleftharpoons}} Q_n^*, \qquad Q_n^* + S_s^* \underset{k_4}{\overset{k_3}{\rightleftharpoons}} C_{n+1,s}^* + E^*, \qquad n = 1, 2, \ldots, N-1. \tag{S1b}$$

The subscript $s$ denotes surface concentration and stars denote dimensional variables. Concentration fields are summarised in Table S1 and rate constants in Table S2; the tables include related dimensionless quantities that will emerge later in this analysis. The reactions take place in a Golgi cisterna (illustrated in Figure 2 of the main text) with the enzyme and complex confined to the membrane bounding the cisterna, so that reactions (S1b) take place on this surface with rates $k_1, k_2, k_3, k_4$. The cargo and solute move in the lumen via diffusion, and adsorb onto (and desorb from) the membrane with rates $k_{Ca}$, $k_{Cd}$, $k_{Sa}$, $k_{Sd}$ via (S1a). The cargo polymerizes up to some size $N$.

We assume the cisterna domain $\Omega$ is pancake-shaped, being thin in the $z^*$ direction, with thickness $h^*(x^*, y^*) \equiv h^*(\mathbf{x}_\perp^*)$, relative to the longer $x^*$ and $y^*$ directions, which span the cisterna footprint $\Omega_\perp$. The cisternal volume $|\Omega|$ and footprint area $|\Omega_\perp|$ define a mean thickness $h_0 = |\Omega|/|\Omega_\perp|$ and mean radius $L_0 = \sqrt{|\Omega_\perp|/\pi}$. We assume symmetry about the midplane $z^* = 0$ of the cisterna. Surface concentrations are functions of $\mathbf{x}_\perp^* \in \Omega_\perp$ and time $t^*$; bulk concentrations are functions of $\mathbf{x}^* \in \Omega$ and $t^*$. We consider the limit $h_0/L_0 \ll 1$ ($|\Omega|^2 \ll |\Omega_\perp|^3$) throughout but allow cisternal shape to vary within this constraint, by allowing $h^*$ to be spatially non-uniform, with $\int_{\Omega_\perp} h^*(\mathbf{x}_\perp^*)\,\mathrm{d}\mathbf{x}_\perp^* = |\Omega|$. Geometric quantities are summarised in Table S3.

At $t^* = 0$, cargo $C_1^*$ at uniform concentration $C_b$ and substrate at uniform concentration $S_b$ are introduced into the cisterna in the presence of membrane-bound enzyme. Polymerization then

Table S1: **Concentration fields:** variables in the left-hand column are dimensional; the right-hand columns show dimensionless surface concentrations. Gaps in the right-hand columns reflect reduction of model complexity via thin-layer and continuous approximations, given in Sections S2 and S2.2 respectively.

| Dimensional | | Thin-layer | Continuous |
|---|---|---|---|
| $C_n^*(\mathbf{x}^*, t^*)$ | Bulk polymer concentration | | |
| $C_{n,s}^*(\mathbf{x}_\perp^*, t^*)$ | Surface polymer concentration | $C_n(\mathbf{x}_\perp, t)$ | $\tilde{C}(\nu, \mathbf{x}_\perp, \tilde{t})$ |
| $S^*(\mathbf{x}^*, t^*)$ | Bulk substrate concentration | | |
| $S_s^*(\mathbf{x}_\perp^*, t^*)$ | Surface substrate concentration | $S(\mathbf{x}_\perp, t)$ | |
| $E^*(\mathbf{x}_\perp^*, t^*)$ | Enzyme concentration | $E(\mathbf{x}_\perp, t)$ | $\tilde{E}(\mathbf{x}_\perp, \tilde{t})$ |
| $Q_n^*(\mathbf{x}_\perp^*, t^*)$ | Complex concentration | $Q_n(\mathbf{x}_\perp, t)$ | $\tilde{Q}(\nu, \mathbf{x}_\perp, \tilde{t})$ |
| $\mathcal{M}_\phi^*(t^*)$ | Mass of functional polymer | | |
| $T_{r,50}^*$ | Time at 50% of maximal output | | |
| $\mathcal{P}_{50}^*$ | Production rate | | |

Table S2: **Rate constants**: the left-hand column shows dimensional parameters, those in the right-hand columns are dimensionless (used in the thin-layer and continuous models of Sections S2 and S2.2).

| Dimensional | Rate | Thin-layer | Continuous |
|---|---|---|---|
| $k_{Sa}$ | Substrate adsorption | | |
| $k_{Sd}$ | Substrate desorption | | $\alpha_S = \frac{k_{Sd}\lvert\Omega\rvert}{2k_{Sa}\lvert\Omega_\perp\rvert}$ |
| $k_{Ca}$ | Polymer adsorption | | |
| $k_{Cd}$ | Polymer desorption | $\alpha_C = \frac{k_{Cd}\lvert\Omega\rvert}{2k_{Ca}\lvert\Omega_\perp\rvert}$ | $\alpha_C$ |
| $k_1$ | Complex formation | | |
| $k_2$ | Complex dissociation | $K_2 = \frac{k_2}{k_1\mathcal{C}} \frac{2k_{Ca}\lvert\Omega_\perp\rvert + k_{Cd}\lvert\Omega\rvert}{k_{Ca}}$ | $K_2$ |
| $k_3$ | Product formation | $K_3 = k_3/k_1$ | $\beta$ |
| $k_4$ | Product dissociation | $K_4 = k_4/k_1$ | $K_4$ |

Table S3: **Geometric quantities**: dimensional (Dim.) input parameters are given in the left-hand column. Dimensionless parameters and variables are given in the right-hand column.

| Dim. | | | Dimensionless |
|---|---|---|---|
| $\lvert\Omega\rvert$ | | Lumen volume | |
| $\lvert\Omega_\perp\rvert$ | | Cisternal footprint area | |
| | $h_0$ | Mean thickness $\lvert\Omega\rvert/\lvert\Omega_\perp\rvert$ | Aspect ratio $\sqrt{\pi}\lvert\Omega\rvert/\lvert\Omega_\perp\rvert^{3/2}$ |
| | $L_0$ | Mean radius $\sqrt{\lvert\Omega_\perp\rvert/\pi}$ | $\mathbf{x}_\perp = \mathbf{x}_\perp^*/L_0$ |
| | $h^*(\mathbf{x}_\perp^*)$ | Height profile | $h(\mathbf{x}_\perp)$ |

Table S4: **Biochemical and transport parameters**: dimensionless parameters are given in the right-hand columns. The left-hand column gives dimensional (Dim.) input parameters to the model. The right-hand columns give parameters used in the thin-layer and continuous models given in Sections S2 and S2.2. "Early" concentrations are those arising after the initial concentration field has equilibrated via adsorption to interfaces. The bottom row shows how time is measured in the thin-layer and continuous models.

| Dim. | | | Thin-layer | Continuous |
|---|---|---|---|---|
| $N$ | | Maximum polymer length | $N$ | |
| $\mathcal{C}$ | | Initial monomer mass | | |
| | $C_b$ | Initial bulk monomer concentration $\mathcal{C}/\|\Omega\|$ | | |
| | $C_0$ | Early surface monomer concentration $\frac{C_b h_0}{2(1+\alpha_C)}$ | | |
| $\mathcal{S}$ | | Initial substrate mass | | |
| | $S_b$ | Initial bulk substrate concentration $\mathcal{S}/\|\Omega\|$ | | |
| | $S_0$ | Early surface substrate concentration $\frac{S_b h_0}{2(1+\alpha_S)}$ | $\sigma = \frac{k_{Sa} S_b (2k_{Ca}\|\Omega_\perp\| + k_{Cd}\|\Omega\|)}{k_{Ca} C_b (2k_{Sa}\|\Omega_\perp\| + k_{Sd}\|\Omega\|)}$ | |
| $\mathcal{E}$ | | Total enzyme mass | $\varepsilon = \frac{\mathcal{E}(2k_{Ca}\|\Omega_\perp\| + k_{Cd}\|\Omega\|)}{2k_{Ca} C_b \|\Omega\|\|\Omega_\perp\|}$ | |
| | $E_0$ | Mean enzyme concentration $\mathcal{E}/(2\|\Omega_\perp\|)$ | | |
| $\phi$ | | Functional polymer threshold | $\phi$ | $\phi$ |
| $D_C$ | | Monomer/polymer diffusivity | $\delta_C = \pi D_C/(k_1\mathcal{E})$ | $\mathcal{D}$ |
| $D_S$ | | Substrate diffusivity | $\delta_S = \pi D_S/(k_1\mathcal{E})$ | |
| $t^*$ | | Time | $t = k_1\mathcal{E}t^*/(2\|\Omega_\perp\|)$ | $\tilde{t} = \frac{t}{N^2(K_2+\sigma K_3)}$ |

takes place until the contents are released (at some time $t^* = T_r^*$). Thus the initial total masses of polymer, substrate and enzyme are $\mathcal{C} = C_b|\Omega|$, $\mathcal{S} = S_b|\Omega|$ and $\mathcal{E}$ respectively, as summarised in Table S4. We will treat $\mathcal{C}$, $\mathcal{S}$ and $\mathcal{E}$ as primary input parameters. To accommodate heterogeneity, the initial enzyme distribution $E^*(\mathbf{x}_\perp^*, 0)$ is given in terms of a dimensionless function $F_E(\mathbf{x}_\perp^*/L_0)$ satisfying $\int_{\Omega_\perp} F_E(\mathbf{x}_\perp^*/L_0)\,\mathrm{d}\mathbf{x}_\perp^* = |\Omega_\perp|$, via $E^*(\mathbf{x}_\perp^*, 0) = E_0 F_E(\mathbf{x}_\perp^*/L_0)$ where $E_0 = \mathcal{E}/(2|\Omega_\perp|)$. Thus

$$\mathcal{C} = \int_\Omega C_1^*(\mathbf{x}^*, 0)\,\mathrm{d}\mathbf{x}^* = C_b|\Omega|, \quad \mathcal{S} = \int_\Omega S^*(\mathbf{x}^*, 0)\,\mathrm{d}\mathbf{x}^* = S_b|\Omega|, \quad \mathcal{E} = 2\int_{\Omega_\perp} E^*(\mathbf{x}_\perp^*, 0)\,\mathrm{d}\mathbf{x}_\perp^*. \quad \text{(S2)}$$

Parameters describing initial concentration fields are summarised in Table S4.

There are three conservation laws associated with (S1), for enzyme, substrate and cargo in $t^* \geq 0$:

$$2|\Omega|_\perp \left[E^*(\mathbf{x}_\perp^*, t^*) + \textstyle\sum_{n=1}^{N-1} Q_n^*(\mathbf{x}_\perp^*, t^*)\right] = \mathcal{E} F_E(\mathbf{x}_\perp^*/L_0), \quad \text{(S3a)}$$

$$\int_\Omega \left(S^*(\mathbf{x}^*, t^*) + \textstyle\sum_{n=2}^{N}(n-1)C_n^*\right)\mathrm{d}\mathbf{x}^* + 2\int_{\Omega_\perp}\left(S_s^*(\mathbf{x}_\perp^*, t^*) + \textstyle\sum_{n=2}^{N}(n-1)C_{n,s}^* + \sum_{n=2}^{N-1}(n-1)Q_n^*\right)\mathrm{d}\mathbf{x}_\perp^* = \mathcal{S}, \quad \text{(S3b)}$$

$$\int_\Omega \textstyle\sum_{n=1}^{N} C_n^*(\mathbf{x}^*, t^*)\,\mathrm{d}\mathbf{x}^* + 2\int_{\Omega_\perp}\left(\sum_{n=1}^{N-1}(C_{n,s}^* + Q_n^*) + C_{N,s}^*\right)\mathrm{d}\mathbf{x}_\perp^* = \mathcal{C}. \quad \text{(S3c)}$$

Eq. (S3a) is a local constraint because the enyzme and complex are immobile. We assess the output of (S1) at the release time using the mass of initial monomer that has been converted to what we term functional luminal polymer (with chain length $n$ above $\phi N$ for some $\phi \in (0,1)$ for which $\phi N$ is an integer), namely

$$\mathcal{M}^*_\phi(T^*_r) = \int_\Omega \textstyle\sum_{n=\phi N}^{N} C^*_n(\mathbf{x}^*, T^*_r)\,\mathrm{d}\mathbf{x}^*. \tag{S4}$$

The cisterna 'matures' as it passes through the *cis*, *medial* and *trans* regions of the Golgi, before its contents are released. A key quantity of interest is the production rate of functional luminal polymer, which may be influenced by cisternal shape. If the forward reaction rates are sufficiently large compared to the reverse reaction rates, and if $T^*_r$ is sufficiently long, then $\mathcal{M}^*_\phi(T^*_r)$ saturates at its maximum possible value $\mathcal{M}^*_{\phi\,\mathrm{max}}$. This is represented by the first term in (S3c) at late times, when the system comes into equilibrium with $C^*_n$ close to zero except for near $n = N$; the offset from $\mathcal{C}$, described by the second integral in (S3c), corresponds to polymerised substrate that remains adsorbed on cisternal membranes at the release time. We define $T^*_{r,50}$ to be the time needed to achieve 50% of the maximum functional output. We will use the production rate at $T^*_{r,50}$, defined as

$$\mathcal{P}^*_{50} = \frac{\mathcal{M}^*_\phi(T^*_{r,50})}{T^*_{r,50}} \quad \text{where} \quad \mathcal{M}^*_\phi(T^*_{r,50}) = \tfrac{1}{2}\mathcal{M}^*_{\phi,\mathrm{max}}, \tag{S5}$$

as a measure of the overall functional performance of the cisterna, seeking in particular conditions under which this metric might be maximised. In instances where the forward reaction cannot proceed, we set $\mathcal{P}^*_{50} = 0$.

##### S1.1 Combining bulk transport with surface reaction

The bulk concentration $C^*(\mathbf{x}^*, t^*)$ of a solute (such as $C^*_n$) in the cisterna is assumed to evolve via diffusion according to

$$C^*_{t^*} = D_C \nabla^{*2} C^* \quad (\mathbf{x}^* \in \Omega), \tag{S6}$$

for some diffusivity $D_C$, where a subscript $t^*$ denotes a time derivative. (We proceed in this Section by considering a single species, before generalising to the full system in Section S1.2.) The concentration averaged across the depth of the cisterna is

$$\overline{C^*}(\mathbf{x}^*_\perp, t^*) = \frac{1}{h^*}\int_{-h^*/2}^{h^*/2} C^*(\mathbf{x}^*_\perp, z^*, t^*)\,\mathrm{d}z^*. \tag{S7}$$

For a thin domain, we can write $C^* = \overline{C^*} + \hat{C}^*$ where the magnitude of the $z^*$-dependent perturbation field $|\hat{C}^*|$ is regulated by the strength of $z^*$-diffusion relative to any reaction. Substitution into (S6) and integration across the cisterna, noting that $\overline{\hat{C}^*} \equiv 0$ and that $|\hat{C}^*| \ll |\overline{C^*}|$ for a thin domain $\Omega$ with $h_0 \ll L_0$, shows that the cross-cisterna average satisfies

$$h^* \overline{C^*}_{t^*} = D_C \nabla^*_\perp \cdot (h^* \nabla^*_\perp \overline{C^*}) - 2 D_C C^*_{\mathsf{n}}|_{\partial\Omega}. \tag{S8}$$

Here $\mathsf{n}$ is the normal coordinate to the cisterna boundary $\partial\Omega$ pointing into the lumen (Fig. 2 of the main text); $C^*_{\mathsf{n}}|_{\partial\Omega}$ is the directional derivative of the bulk field evaluated at the surface. Eq. (S8) uses the thin-layer approximation $C^*_{\mathsf{n}} \approx \mp\left(\hat{C}^*_{z^*} - \tfrac{1}{2}\nabla^*_\perp h^* \cdot \nabla^*_\perp \overline{C^*}\right)$ on $z^* = \pm\tfrac{1}{2}h^*$, as exploited for example in Ref. 47 in the main text. Balancing terms in this expression suggests that $\hat{C}^*$ is a factor $(h_0/L_0)^2$ smaller than $\overline{C^*}$, making the error in (S8) of $O((h_0/L_0)^2)$.

The flux between bulk and surface, such as in (S1a), is modelled by linear kinetics and is assumed to be mediated by bulk diffusion according to

$$k_{Ca} C^*|_{\partial\Omega} - k_{Cd} C_s^* = D_C C_{\mathsf{n}}^* \quad (\mathbf{x}^* \in \partial\Omega). \tag{S9}$$

Writing $C^*|_{\partial\Omega} \approx \overline{C^*}$, we can then combine the evolution equation for surface concentration, $C_{s,t^*}^* = f(C_s^*, \dots) + D_C C_{\mathsf{n}}^*$, for some surface reaction term $f$, with the bulk transport equation (S8) to give the coupled system

$$C_{s,t^*}^* = f(C_s^*, \dots) + k_{Ca} C^*|_{\partial\Omega} - k_{Cd} C_s^* \tag{S10a}$$

$$h^* \overline{C}_{t^*}^* = D_C \nabla_\perp^* \cdot (h^* \nabla_\perp^* \overline{C^*}) - 2(k_{Ca} C^*|_{\partial\Omega} - k_{Cd} C_s^*). \tag{S10b}$$

Adding twice (S10a) to (S10b), it follows that

$$h^* \overline{C^*}_{t^*} = D_C \nabla_\perp^* \cdot (h^* \nabla_\perp^* \overline{C^*}) - 2[C_{s,t^*}^* - f(C_s^*, \dots)]. \tag{S11}$$

If the exchange between bulk and surface is very rapid, the surface and bulk concentrations quickly come into equilibrium and are related (balancing adsorption and desorption terms in (S10)) by

$$\overline{C^*}(\mathbf{x}_\perp^*, t^*) \approx C^*|_{\partial\Omega}(\mathbf{x}_\perp^*, t^*) \approx \frac{k_{Cd}}{k_{Ca}} C_s^*(\mathbf{x}_\perp^*, t^*). \tag{S12}$$

The factor $k_{Cd}/k_{Ca}$ has dimensions of inverse length, converting a surface concentration into a bulk concentration. In the strong-exchange, thin-film limit (S12), (S11) can be written as an evolution equation for the surface concentration

$$\left[1 + \frac{h^* k_{Cd}}{2k_{Ca}}\right] C_{s,t^*}^* = f(C_s^*, \dots) + \frac{k_{Cd}}{2k_{Ca}} D_C \nabla_\perp^* \cdot (h^* \nabla_\perp^* C_s^*). \tag{S13}$$

Although the reaction takes place on the surface, (S13) captures the fact that the solute may spend significant time in the bulk, where it can diffuse laterally. The kinetics are mediated by the affinity of the solute for the membrane. This is regulated by the ratio of local lumen thickness $h^*(\mathbf{x}_\perp)$ to the adsorption depth $h_C \equiv 2k_{Ca}/k_{Cd}$. Given $C_s^*(\mathbf{x}_\perp^*, t^*)$, the total amount of solute in the cisterna can be written

$$\int_\Omega C^* \, \mathrm{d}\mathbf{x}^* + 2 \int_{\Omega_\perp} C_s^* \, \mathrm{d}\mathbf{x}_\perp^* = \int_{\Omega_\perp} \left(2 + \frac{h^* k_{Cd}}{k_{Ca}}\right) C_s^*(\mathbf{x}_\perp^*, t) \, \mathrm{d}\mathbf{x}_\perp^*. \tag{S14}$$

To accompany (S13), we impose no-flux conditions on the periphery $\partial\Omega_\perp$ of $\Omega_\perp$

$$\mathbf{n}_\perp \cdot (h^* \nabla_\perp^* C_s^*) = 0, \quad (\mathbf{x}_\perp^* \in \partial\Omega_\perp), \tag{S15}$$

where $\mathbf{n}_\perp$ is the normal to $\partial\Omega_\perp$ in the plane $z^* = 0$.

#### S1.2 Combining transport and kinetics

Using the reaction-diffusion formalism (S13), assuming the cisterna is thin and bulk-surface exchange is rapid for both substrate and polymer, we can now write evolution equations for the system (S1) in terms of surface concentrations $C^*_{n,s}(\mathbf{x}^*_\perp, t^*)$, $E^*(\mathbf{x}^*_\perp, t^*)$, $Q^*_n(\mathbf{x}^*_\perp, t^*)$ and $S^*_s(\mathbf{x}^*_\perp, t^*)$. These are

$$\left[1+\frac{h^* k_{Cd}}{2k_{Ca}}\right] C^*_{n,s,t^*} = -k_1 C^*_{n,s}E^*(1-\delta_{nN}) + k_2 Q^*_n + k_3 Q^*_{n-1}S^*_s - k_4 C^*_{n,s}E^*(1-\delta_{n1}) + \frac{k_{Cd}}{2k_{Ca}} D_C \nabla^*_\perp \cdot (h^* \nabla_\perp C^*_{n,s}), \qquad (n=1,\ldots,N) \tag{S16a}$$

$$E^*_{t^*} = \textstyle\sum_{n=1}^N \left[-k_1 C^*_{n,s}E^*(1-\delta_{nN}) + k_2 Q^*_n + k_3 Q^*_n S^*_s - k_4 C^*_{n,s}E^*(1-\delta_{n1})\right], \tag{S16b}$$

$$Q^*_{n,t} = k_1 C^*_{n,s}E^* - k_2 Q^*_n - k_3 Q^*_n S^*_s + k_4 C^*_{n+1,s}E^*, \qquad (n=1,\ldots,N-1) \tag{S16c}$$

$$\left[1+\frac{h^* k_{Sd}}{2k_{Sa}}\right] S^*_{t^*} = \textstyle\sum_{n=1}^{N-1}\left[-k_3 Q^*_n S^*_s + k_4 C^*_{n+1,s}E^*\right] + \frac{k_{Sd}}{2k_{Sa}} D_S \nabla^*_\perp \cdot (h^* \nabla^*_\perp S^*_s). \tag{S16d}$$

We enforce $Q^*_0 \equiv 0$ and $Q^*_N \equiv 0$ and use $\delta$-functions to switch off the terms $k_4 C^*_{1,s}E^*$ and $k_1 C^*_{N,s}E^*$ in (S16a,b). $E^*$ and $Q^*_n$ remain bound to the membrane, so cannot be transported in the bulk. There is competition among all species for enzyme and substrate. $D_C$ and $D_S$ are bulk diffusivities; we assume that a single diffusivity can be used for all polymer sizes, which is reasonable provided the overall reaction is not diffusion-limited. Imposing no-flux boundary conditions on $\partial\Omega_\perp$, it can be verified that (S16) satisfies

$$E^*_{t^*} + \textstyle\sum_{n=1}^{N-1} Q^*_{n,t^*} = 0, \tag{S17a}$$

$$\int_{\Omega_\perp}\left(\left[1+\frac{h^* k_{Sd}}{2k_{Sa}}\right] S^*_{t^*} + \left[1+\frac{h^* k_{Cd}}{2k_{Ca}}\right] \textstyle\sum_{n=2}^{N}(n-1) C^*_{n,s,t^*} + \textstyle\sum_{n=2}^{N-1}(n-1) Q^*_{n,t^*}\right) \mathrm{d}\mathbf{x}^*_\perp = 0, \tag{S17b}$$

$$\int_{\Omega_\perp}\left(\left[1+\frac{h^* k_{Cd}}{2k_{Ca}}\right] \textstyle\sum_{n=1}^{N} C^*_{n,s,t} + \textstyle\sum_{n=1}^{N-1} Q^*_{n,t^*}\right) \mathrm{d}\mathbf{x}^*_\perp = 0, \tag{S17c}$$

consistent with the conservation laws (S3).

Let $T_0 = 1/(k_1 E_0)$ be a reference reaction timescale. Then we can define dimensionless diffusivities and capacitances as

$$\delta_C = \frac{D_C}{2L_0^2 k_1 E_0}, \quad \delta_S = \frac{D_S}{2L_0^2 k_1 E_0}, \quad \alpha_C = \frac{k_{Cd}|\Omega|}{2k_{Ca}|\Omega_\perp|}, \quad \alpha_S = \frac{k_{Sd}|\Omega|}{2k_{Sa}|\Omega_\perp|} \tag{S18}$$

(see Tables S2 and S4 for definitions of these quantities in terms of input parameters). Factors of 2 are included to account for the fact that reactions take place on two membranes. $\delta_C$ ($\delta_S$) measures a reaction time versus the time for the cargo (substrate) to diffuse along the length of the cisterna. $\alpha_C$ ($\alpha_S$) measures the solubility of the cargo (substrate); equivalently, $1/\alpha_C$ measures the cargo's affinity for the membrane. When $\alpha_C \gg 1$ in (S18), so that the adsorption depth $h_C = 2k_{Ca}/k_{Cd}$ is much smaller than the mean cisternal thickness $h_0$, the cargo adsorbs weakly onto the interface and temporal evolution of the concentration field is strongly mediated by bulk transport, and hence by the shape of the domain via the prefactor $h^*$ of $C^*_{s,t^*}$ in (S16a).

We assume that the initial conditions (S2) (and those implicit in (S3)) equilibrate rapidly, via (S12) and (S14), to the form

$$C^*_{n,s}(\mathbf{x}^*_\perp,0)=C_0F_n(\mathbf{x}^*_\perp/L_0),\quad S^*_s(\mathbf{x}^*_\perp,0)=S_0,\quad E^*(\mathbf{x}^*_\perp,0)=E_0F_E(\mathbf{x}^*_\perp/L_0),\quad Q^*_n(\mathbf{x}^*_\perp)=0, \tag{S19}$$

for some dimensionless functions $F_n$, where $F_1=1$ and $F_n=0$ for $n=2,\ldots,N$. $C_0$ and $S_0$ are the surface concentrations that form rapidly after the system is initialised in the bulk (via (S13)), satisfying

$$\mathcal{C}=2C_0|\Omega_\perp|(1+\alpha_C),\quad \mathcal{S}=2S_0|\Omega_\perp|\left(1+\alpha_S\right),\quad \mathcal{E}=2E_0|\Omega_\perp|. \tag{S20}$$

### S2 The nondimensional thin-layer model

As a first step in simplifying the model, we nondimensionalise (S16) and (S19) by setting

$$\mathbf{x}^*_\perp=L_0\mathbf{x}_\perp,\quad h^*(\mathbf{x}^*_\perp)=h_0h(\mathbf{x}_\perp)\quad C^*_{n,s}(\mathbf{x}^*_\perp,t^*)=C_0C_n(\mathbf{x}_\perp,t),\quad Q^*_n(\mathbf{x}^*_\perp,t^*)=E_0Q_n(\mathbf{x}_\perp,t), \tag{S21a}$$

$$t^*=T_0t\qquad T^*_r=T_0T_r,\qquad E^*(\mathbf{x}^*_\perp,t^*)=E_0E(\mathbf{x}_\perp,t),\qquad S^*_s(\mathbf{x}^*_\perp,t^*)=S_0S(\mathbf{x}_\perp,t). \tag{S21b}$$

Subscripts $s$ have been removed from dimensionless surface concentrations; dimensionless variables are summarised in Table S1. We define dimensionless concentration ratios and rate constants as

$$\varepsilon=\frac{E_0}{C_0}=\frac{\mathcal{E}(1+\alpha_C)}{\mathcal{C}},\quad \sigma=\frac{S_0}{C_0}=\frac{\mathcal{S}(1+\alpha_C)}{\mathcal{C}(1+\alpha_S)},\quad K_2=\frac{k_2}{k_1C_0},\quad K_3=\frac{k_3}{k_1},\quad K_4=\frac{k_4}{k_1} \tag{S22}$$

(see Tables S2 and S4). In dimensionless variables, the lumen area and volume become

$$\int_{\Omega_\perp}\mathrm{d}\mathbf{x}_\perp=\pi,\quad \int_{\Omega_\perp}h(\mathbf{x}_\perp)\,\mathrm{d}\mathbf{x}_\perp=\pi \tag{S23}$$

and (S16) is

$$\begin{aligned}[1+\alpha_Ch]\,C_{n,t}&=-C_nE(1-\delta_{nN})+K_2Q_n+K_3\sigma Q_{n-1}S-K_4C_nE(1-\delta_{n1})\\&\quad+\alpha_C\delta_C\nabla_\perp\cdot(h\nabla_\perp C_n),\qquad (n=1,\ldots,N),\end{aligned} \tag{S24a}$$

$$\varepsilon E_t=\textstyle\sum_{n=1}^{N}\left[-C_nE(1-\delta_{nN})+K_2Q_n+K_3\sigma Q_nS-K_4C_nE(1-\delta_{n1})\right], \tag{S24b}$$

$$\varepsilon Q_{n,t}=C_nE-K_2Q_n-K_3\sigma Q_nS+K_4C_{n+1}E,\qquad (n=1,\ldots,N-1), \tag{S24c}$$

$$[1+\alpha_Sh]\,S_t=\textstyle\sum_{n=1}^{N-1}\left[-K_3Q_nS+(K_4/\sigma)C_{n+1}E\right]+\alpha_S\delta_S\nabla_\perp\cdot(h\nabla_\perp S), \tag{S24d}$$

with initial conditions (S19) becoming

$$C_n(\mathbf{x}_\perp,0)=F_n(\mathbf{x}_\perp),\quad E(\mathbf{x}_\perp,0)=F_E(\mathbf{x}_\perp),\quad Q_n(\mathbf{x}_\perp,0)=0,\quad S(\mathbf{x}_\perp,0)=1. \tag{S24e}$$

Imposing no-flux conditions on $C_n$ and $S$ at $\partial\Omega_\perp$, the conservation laws (S3) become

$$E(\mathbf{x}_\perp,t)+\textstyle\sum_{n=1}^{N-1}Q_n(\mathbf{x}_\perp,t)=F_E(\mathbf{x}_\perp), \tag{S25a}$$

$$\int_{\Omega_\perp}\Big[(1+h\alpha_S)\,\sigma S+(1+h\alpha_C)\textstyle\sum_{n=2}^{N}(n-1)C_n+\varepsilon\sum_{n=2}^{N-1}(n-1)Q_n\Big]\,\mathrm{d}\mathbf{x}_\perp=\pi\sigma(1+\alpha_S), \tag{S25b}$$

$$\int_{\Omega_\perp}\Big[(1+h\alpha_C)\textstyle\sum_{n=1}^{N}C_n+\varepsilon\sum_{n=1}^{N-1}Q_n\Big]\,\mathrm{d}\mathbf{x}_\perp=\pi(1+\alpha_C), \tag{S25c}$$

where $\int_{\Omega_\perp} F_E(\mathbf{x}_\perp)\,\mathrm{d}\mathbf{x}_\perp = \pi$. The dimensional bulk mass of functional polymer (S4) is (using (S12), (S20) and (S21))

$$\mathcal{M}^*_\phi(T^*_r) = \frac{\mathcal{C}\alpha_C}{\pi(1+\alpha_C)} \int_{\Omega_\perp} \textstyle\sum_{n=\phi N}^{N} h(\mathbf{x}_\perp) C_n(\mathbf{x}_\perp, T_r)\,\mathrm{d}\mathbf{x}_\perp. \tag{S26}$$

Of the total mass of substrate $\mathcal{C}$, the proportion in the bulk at large times, once the reaction has pushed polymers towards the maximum length $N$, is given by the ratio of the second to the fourth and fifth term in (S25c), explaining the factor $\alpha_C/(\pi(1+\alpha_C))$ in (S26). Simulation can in principle be used to identify the large-time limit $\mathcal{M}^*_{\phi\,\max}$, the time $T^*_{r,50}$ at which $\mathcal{M}^*_\phi = \mathcal{M}^*_{\phi\,\max}/2$ and hence $\mathcal{P}^*_{50}$ via (S5), which becomes

$$\mathcal{P}^*_{50}(T^*_{r,50}) \equiv \frac{\mathcal{M}^*_\phi(T^*_{r,50})}{T^*_{r,50}} = \frac{\mathcal{C}\alpha_C}{1+\alpha_C}\cdot\frac{k_1\mathcal{E}}{2\pi|\Omega_\perp|}\cdot\frac{1}{T_{r,50}}\int_{\Omega_\perp} \textstyle\sum_{n=\phi N}^{N} h(\mathbf{x}_\perp) C_n(\mathbf{x}_\perp, T_{r,50})\,\mathrm{d}\mathbf{x}_\perp. \tag{S27}$$

The production rate is proportional to $k_1\mathcal{C}\mathcal{E}/|\Omega_\perp|$ multiplied by dimensionless quantities. This dimensional prefactor illustrates how a large surface area can slow the production rate by diluting the available enzyme.

The original model (S16) involves 17 parameters (8 rate constants, 2 geometric factors, 7 biochemical and transport parameters in the left-hand columns of Tables S2, S3 and S4 respectively). The nondimensional model (S24) involves 11 parameters (5 rate constants, 6 biochemical and transport parameters identified in columes 3 and 4 of Tables S2 and S4 respectively). We now simplify the model further, reducing the number of independent input parameters to 6, in order to obtain a simpler prediction of the production rate $\mathcal{P}^*_{50}$.

#### S2.1 Model reduction: limited enzyme and abundant substrate

The system (S24) involves the evolution of $2N+2$ variables. We can reduce this number while retaining essential features of the dynamics, as follows.

Suppose that there is limited enzyme, so that $\varepsilon \ll 1$ (see (S22)). After an initial adjustment over a timescale $O(\varepsilon)$, $Q_n$ and $E$ come into equilibrium with the concentrations $C_n$ so that (S24c) and (S25a) give

$$(K_2 + \sigma S K_3) Q_n = (C_n + K_4 C_{n+1}) E, \quad E = F_E(\mathbf{x}_\perp) - \textstyle\sum_{n=1}^{N-1} Q_n. \tag{S28}$$

We can use (S25a) instead of (S24b) to specify $E$. We then use (S28) to write (S24a,d,e] as

$$\begin{aligned}[1+\alpha_C h]\, C_{n,t} = K_3\sigma S\,[Q_{n-1}(1-\delta_{n1}) - Q_n(1-\delta_{nN})] + K_4 E\,[C_{n+1}(1-\delta_{nN}) - C_n(1-\delta_{n1})] \\ + \alpha_C\delta_C\nabla_\perp\cdot(h\nabla_\perp C_n), \qquad C_n(\mathbf{x}_\perp,0) = F_n(\mathbf{x}_\perp),\end{aligned} \tag{S29a}$$

$$[1+\alpha_S h]\, S_t = \textstyle\sum_{n=1}^{N-1}\left[-K_3 Q_n S + (K_4/\sigma) C_{n+1} E + \alpha_S\delta_S\nabla_\perp\cdot(h\nabla_\perp S)\right], \qquad S(\mathbf{x}_\perp,0) = 1, \tag{S29b}$$

which is an $(N+1)$th-order system.

Suppose also that the substrate is abunbdant, so that $\sigma \gg 1$ (see (S22)), and that $K_3 \sim (1/\sigma) \ll 1 \sim K_4$, where $\sim$ denotes 'scales like' (in the limit $\sigma \gg 1$). Then the dominant problem for $S$, (S29b), is $[1+\alpha_S h]\, S_t = \alpha_S\delta_S\nabla_\perp\cdot(h\nabla_\perp S)$ with $S(\mathbf{x}_\perp,0) = 1$, implying that $S$ remains at unity to leading order. This reduces the problem to (S29a) for $C_n$ plus (S28) for $Q_n$ with $S = 1$, which is an $N$th-order system.

##### S2.2 The continuous model

We proceed assuming that substrate is abundant and enzyme is scarce ($\sigma \gg 1$, $\varepsilon \ll 1$, with $\sigma K_3 \sim K_4 \sim 1$ in the limit) and therefore rate-limiting. Equations (S24, S25) become

$$
\begin{aligned}
(1+\alpha_C h)C_{n,t} &= -C_n E(1-\delta_{nN}) + K_2 Q_n + K_3\sigma Q_{n-1} - K_4 C_n E(1-\delta_{n1}) \\
&\qquad + \alpha_C\delta_C \nabla_\perp\cdot(h\nabla_\perp C_n), \qquad (n=1,2,\ldots,N) \quad \text{(S30a)}\\
0 &= C_n E - K_2 Q_n - K_3\sigma Q_n + K_4 C_{n+1}E, \qquad (n=1,2,\ldots N-1), \quad \text{(S30b)}
\end{aligned}
$$

$$F_E(\mathbf{x}_\perp) = E + \textstyle\sum_{n=1}^{N-1} Q_n, \qquad \displaystyle\int_{\Omega_\perp} F_E(\mathbf{x}_\perp)\,\mathrm{d}\mathbf{x}_\perp = \pi, \tag{S30c}$$

$$\pi(1+\alpha_C) = \int_{\Omega_\perp}\left[(1+h(\mathbf{x}_\perp)\alpha_C)\textstyle\sum_{n=1}^{N} C_n(\mathbf{x}_\perp,t)\right]\mathrm{d}\mathbf{x}_\perp, \tag{S30d}$$

with which we wish to determine the production rate via (S26, S27). At large times, diffusion will eliminate spatial gradients in $C_n$ and (S23) applied to (S30d) shows that $\sum_{n=1}^{\infty} C_n \to 1$ as $t\to\infty$. For a strong forward region, $\sum_{n=\phi N}^{N} C_n \to 1$ in this limit, making $\mathcal{M}^*_{\phi\,\max} = \alpha_C\mathcal{C}/(1+\alpha_C)$. Since $\alpha_C = h_0/h_C$ (see (S18)), where $h_0 \equiv |\Omega|/|\Omega_\perp|$ and $h_C \equiv 2k_{Ca}/k_{Cd}$, the maximum polymer production increases with mean cisternal depth $h_0$, but only over the range in which $h_0$ is comparable to the polymer adsorption depth $h_C$.

Eq. (S30) is a hybrid discrete-continuous system. It is helpful to reduce it to a fully continuous system. Taking $N \gg 1$, we introduce $\nu = n/N$, a continuous variable in $(0,1]$ and set

$$C_n(\mathbf{x}_\perp,t) \approx \frac{1}{N}\tilde{C}(\nu,\mathbf{x}_\perp,\tilde{t}), \quad Q_n(\mathbf{x}_\perp,t) \approx \frac{1}{N}\tilde{Q}(\nu,\mathbf{x}_\perp,\tilde{t}), \quad E(\mathbf{x}_\perp,t) = \tilde{E}(\mathbf{x}_\perp,\tilde{t}), \tag{S31}$$

(variables are listed in Table S1), where $\tilde{C}$ and $\tilde{Q}$ are smooth densities that interpolate $C_n$ and $Q_n$ respectively and $\tilde{t}$ will be defined shortly. Then, Taylor-expanding, we have

$$NC_{n\pm1}(t) \approx \tilde{C}(\nu\pm(1/N),\tilde{t}) \approx \tilde{C}(\nu,\tilde{t}) \pm \frac{1}{N}\tilde{C}_\nu(\nu,\tilde{t}) + \frac{1}{2N^2}\tilde{C}_{\nu\nu}(\nu,\tilde{t}) + O(1/N^3). \tag{S32}$$

Away from $\nu=0$ and $\nu=1$, and introducing the continuous concentration field, (S30a) becomes

$$(1+\alpha_C h)\tilde{C}_{\tilde{t}}\frac{\mathrm{d}\tilde{t}}{\mathrm{d}t} = \frac{\tilde{E}}{N}\tilde{C}_\nu\frac{K_2K_4-\sigma K_3}{K_2+\sigma K_3} + \frac{\tilde{E}}{2N^2}\tilde{C}_{\nu\nu}\frac{K_2K_4+\sigma K_3}{K_2+\sigma K_3} + \alpha_C\delta_C\nabla_\perp\cdot(h\nabla_\perp\tilde{C}). \tag{S33}$$

We set $\sigma K_3 = K_2K_4 + \beta/N$, treating $\beta = O(1)$ as a parameter that perturbs the balance between forwards and reverse reactions (so that $\beta > 0$ drives the reaction forwards). In terms of the original variables,

$$
\begin{aligned}
\beta &= \frac{N(2k_{Ca}|\Omega_\perp| + k_{C_d}|\Omega|)}{C_b k_1 k_{Ca}}\left(\frac{S_b k_3 k_{Sa}}{2k_{Sa}|\Omega_\perp| + k_{Sd}|\Omega|} - \frac{k_2k_4}{k_1|\Omega|}\right)\\
&= \frac{N}{\mathcal{C}k_1k_{Ca}}\cdot\frac{2k_{Ca}|\Omega_\perp| + k_{C_d}|\Omega|}{2k_{Sa}|\Omega_\perp| + k_{Sd}|\Omega|}\cdot\frac{(S_bk_1k_3k_{Sa} - k_2k_4k_{Sd})|\Omega| - 2k_2k_4k_{Sa}|\Omega_\perp|}{k_1},
\end{aligned} \tag{S34}
$$

which encapsulates the full set of reactions (S1) (see Table S2). We assume that

$$S_bk_1k_3k_{Sa} > k_2k_4k_{Sd}, \tag{S35}$$

which ensures that there is sufficient substrate for the forward reactions in (S1) to overcome the reverse reactions. Since $S_b = \mathcal{S}/|\Omega|$, (S35) places an upper bound on the cisterna volume, ensuring that the substrate is sufficiently concentrated for polymerisation to proceed. In addition, for $\beta > 0$ (S34b) provides the stronger constraint

$$\frac{|\Omega|}{|\Omega_\perp|} < \frac{2k_{Sa}}{k_{Sd}} \left( \frac{\mathcal{S}k_1 k_3}{2|\Omega_\perp| k_2 k_4} - 1 \right) \equiv h_{\text{cut-off}} \tag{S36}$$

which identifies a maximum cisternal thickness that allows polymerization to proceed. This critical thickness is proportional to the adsorption depth of substrate $h_S \equiv 2k_{Sa}/k_{Sd}$ modified by a factor involving forward over reverse reaction rates, with adsorbed substrate distributed over the two cisternal membranes.

Eq. (S33) is an advection-diffusion equation, with advection taking place in what we may call polymerization space (with coordinate $\nu$) and diffusion taking place in both $\nu$-space and (physical) $\mathbf{x}_\perp$-space. The integral constraints (S30c,d) become

$$F_E(\mathbf{x}_\perp) = \tilde{E} + \int_0^1 \tilde{Q}\, \mathrm{d}\nu, \quad \pi(1+\alpha_C) = \int_{\Omega_\perp} \left[ (1 + h(\mathbf{x}_\perp)\alpha_C) \int_0^1 \tilde{C}(\nu, \mathbf{x}_\perp, \tilde{t})\, \mathrm{d}\nu \right] \mathrm{d}\mathbf{x}_\perp. \tag{S37}$$

To leading order, (S30b) gives $(K_2 + \sigma K_3)\tilde{Q} = \tilde{C}\tilde{E}(1 + K_4)$. Integrating with respect to $\nu$, it follows that the enzyme distribution satisfies

$$\tilde{E}(\mathbf{x}_\perp, \tilde{t}) = F_E(\mathbf{x}_\perp) \left[ 1 + \frac{1 + K_4}{K_2 + \sigma K_3} \int_0^1 \tilde{C}(\nu, \mathbf{x}_\perp, \tilde{t})\, \mathrm{d}\nu \right]^{-1}. \tag{S38}$$

The prefactor of the integral is $1/K_2$ with error $O(1/N)$. Thus, in general, the advection-diffusion system (S33, S38) is non-local, non-linear and inhomogeneous.

To simplify the problem further, we rescale time by $t = \tilde{t}N^2(K_2 + \sigma K_3)$ and assume $\beta = O(1)$ for $N \gg 1$. We have now reached the final reduction of the model, in which we write (S33) as

$$(1 + \alpha_C h)\tilde{C}_{\tilde{t}} + \tilde{E}\beta\tilde{C}_\nu = \tilde{E}K_2K_4\tilde{C}_{\nu\nu} + \mathcal{D}\nabla_\perp \cdot (h\nabla_\perp \tilde{C}), \tag{S39a}$$

$$\tilde{E}(\mathbf{x}_\perp, \tilde{t}) = F_E(\mathbf{x}_\perp) \left[ 1 + \frac{1}{K_2} \int_0^1 \tilde{C}(\nu, \mathbf{x}_\perp, \tilde{t})\, \mathrm{d}\nu \right]^{-1}, \tag{S39b}$$

where $\mathcal{D} = \alpha_C \delta_C N^2(K_2 + \sigma K_3)$. In terms of the original parameters,

$$\mathcal{D} = \frac{k_{Cd}\pi D_C N^2 (2k_{Ca}|\Omega_\perp| + k_{Cd}|\Omega|)\left(2k_2k_{Sa}|\Omega_\perp| + (k_2k_{Sd}|\Omega| + k_3k_{Sa}\mathcal{S})\right)}{2k_{Ca}^2 k_1^2 |\Omega_\perp||\Omega|\mathcal{C}\mathcal{E}(2k_{Sa}|\Omega_\perp| + k_{Sd}|\Omega|)}. \tag{S40}$$

We impose no-flux conditions (in physical space) (S15) on $\partial\Omega_\perp$ plus no-flux conditions (in polymerization space)

$$\beta\tilde{C} = K_2K_4\tilde{C}_\nu \quad \text{at} \quad \nu = 0, 1 \tag{S41}$$

and the (nominal) initial condition

$$\tilde{C}(\nu, \mathbf{x}_\perp, 0) = NH(N^{-1} - \nu), \tag{S42}$$

Table S5: **Parameter combinations relevant to the production rate (S47).**

| | |
|---|---|
| Cargo adsorption thickness | $h_C = 2k_{Ca}/k_{Cd}$ |
| Substrate adsorption thickness | $h_S = 2k_{Sa}/k_{Sd}$ |
| Adsorption thickness ratio | $\lambda = h_C/h_S$ |
| | $\zeta = 2k_2\lvert\Omega_\perp\rvert/(k_3\mathcal{S})$ |
| | $\gamma = 2k_2\lvert\Omega_\perp\rvert/(k_1\mathcal{C})$ |
| | $\Delta = 2k_2k_4\lvert\Omega_\perp\rvert/(k_1k_3\mathcal{S})$ |

where $H$ is the Heaviside function, ensuring that $\int_0^1 \tilde{C}(\nu, \mathbf{x}_\perp, 0)\,\mathrm{d}\nu = 1$. Eq. (S41) ensures that the conservation law (S37b) is satisfied under (S39). Having solved (S39), we integrate over space to determine the final bulk distribution of luminal polymer

$$\tilde{M}(\nu, \tilde{T}_r) \equiv \int_{\Omega_\perp} h(\mathbf{x}_\perp)\tilde{C}(\nu, \mathbf{x}_\perp, \tilde{T}_r)\,\mathrm{d}\mathbf{x}_\perp, \tag{S43}$$

leading to dimensional (bulk) functional mass (see (S26))

$$\mathcal{M}^*_\phi(T^*_r) = \frac{\alpha_C\mathcal{C}}{\pi(1+\alpha_C)}\int_\phi^1 \tilde{M}(\nu, \tilde{T}_r)\,\mathrm{d}\nu. \tag{S44}$$

At very large times, with sufficiently large forward reaction rates, the $\tilde{C}$ field can be expected to have no spatial gradients and the conservation law $(\mathrm{S37})_2$ requires that $\int_\phi^1 \tilde{M}\,\mathrm{d}\nu \to \pi$. Thus $\mathcal{M}^*_{\phi\,\max} = \alpha_C\mathcal{C}/(1+\alpha_C)$ and $\tilde{T}_{r,50}$ is defined by

$$\int_\phi^1 \tilde{M}(\nu, \tilde{T}_{r,50})\,\mathrm{d}\nu = \tfrac{1}{2}\pi. \tag{S45}$$

The dimensional release time is

$$T^*_{r,50} = \frac{2|\Omega_\perp|N^2(K_2+\sigma K_3)}{k_1\mathcal{E}}\tilde{T}_{r,50} \tag{S46}$$

which can be written

$$T^*_{r,50} = \frac{2|\Omega_\perp|N^2}{k_1\mathcal{E}}\left[\frac{2k_2|\Omega_\perp|}{k_1\mathcal{C}}\left(1+\frac{h_0}{h_C}\right)+\frac{k_3\mathcal{S}}{k_1\mathcal{C}}\frac{1+(h_0/h_C)}{1+(h_0/h_S)}\right]\tilde{T}_{r,50}, \tag{S47}$$

in terms of the adsorption depth of polymer, $h_C \equiv 2k_{Ca}/k_{Cd}$, and of substrate, $h_S \equiv 2k_{Sa}/k_{Sd}$. Eq. (S47) indicates how $T^*_{r,50}$ varies with mean cisternal depth $h_0$. The term proportional to $k_2$ in (S47) relates to complex formation; it increases linearly with $h_0$, reflecting the way in which a deeper cisterna holds lower concentrations of polymer, slowing polymerisation. The term proportional to $k_3$ varies more weakly with depth, as varying the depth of the cisterna changes the concentration of substrate (promoting the forward reaction) and complex (promoting the reverse reaction). Table S5 lists these and other parameter combinations that influence the production rate.

Likewise, we can write (S27) as

$$\mathcal{P}^*_{50} = \frac{(k_1\mathcal{C})^2\mathcal{E}}{k_3\mathcal{S}}\frac{1}{4|\Omega_\perp|N^2}\frac{(h_0/h_C)(1+(h_0/h_S))}{(1+(h_0/h_C))^2[1+\zeta(1+(h_0/h_s))]}\frac{1}{\tilde{T}_{r,50}}. \tag{S48}$$

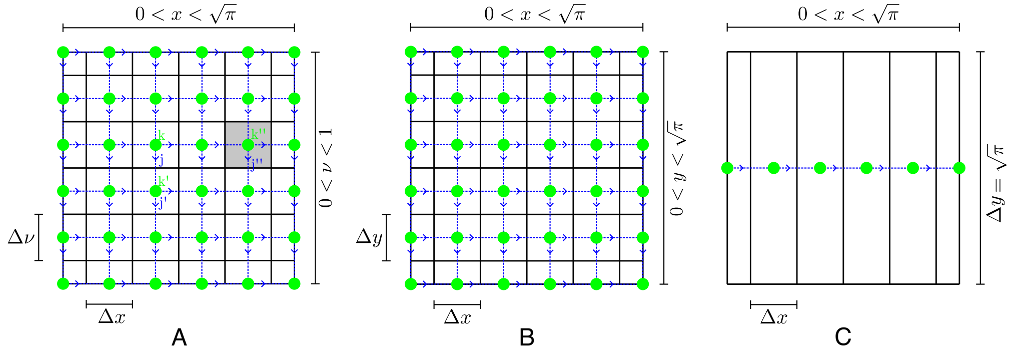


Figure S1: (A) Diagram illustrating discretization of polymerisation space and one spatial dimension. Vertices (green circles labelled by $k$) and surrounding cuboids discretise the domain. Vertices are connected to their immediate neighbours by directed edges (blue dotted lines labelled by $j$). For the labels shown, the incidence matrix has elements $\mathsf{A}_{jk} = -1$, $\mathsf{A}_{jk'} = 1$, $\mathsf{A}_{j'k'} = -1$. The concentration field $\mathsf{C}_k$ is evaluated at vertices and fluxes between vertices are evaluated at edges. (B) Diagram showing discretization of physical space in the case where there is no assumption of uniformity in the second spatial dimension, $y$. This construction is identical to that used for the first spatial dimension $x$ and polymerisation dimension $\nu$. (C) Diagram showing discretization of physical space in the case where solutions are uniform in the second spatial dimension $y$. The construction is similar but uses a single row of vertices in $y$, with $\Delta y$ being the full range of y. All spatial dimensions are normalised to length $\sqrt{\pi}$.

The fraction involving $h_0$ in this expression falls to zero as $h_0 \to 0$ and as $h_0 \to \infty$, suggesting the existence of an optimum cisternal depth that depends on the parameters

$$\zeta = 2k_2|\Omega_\perp|/(k_3\mathcal{S}) \quad \text{and} \quad \lambda = h_C/h_S, \tag{S49}$$

as given in Table S5.

#### S3 Discretization

The discretization scheme used for numerical solutions of (S39) is shown in Fig. S1. An incidence matrix approach is used to construct approximations of the differential operators in this high-dimensional problem. We fill the solution domain, spanning two physical space dimensions and polymerisation space, with point vertices (indexed by $k = 1, \ldots, K$), some of which lie on the boundary of the solution domain, with the remainder at equal intervals within the interior. Each vertex is connected to neighbouring vertices by directed edges (indexed by $j = 1, \ldots, J$). The discretization of the solution domain is defined by cuboids formed from the perpendicular bisecting planes of edges between vertices. With this framework, the system can be described by a signed $J \times K$ incidence matrix $\mathsf{A}$ that maps vertices to edges such that edge $j$ leaving (entering) vertex $k$ has $A_{jk} = -1$ (1). For any edge $j$ and vertex $k$ that are not incident, $A_{jk} = 0$.

The discretized concentration field can be expressed as a vector $\mathsf{C}$ of values over vertices, while discretized flux is evaluated over edges. The cross-sectional capacity of a given edge is equal to the area ($\Delta x \Delta y$, $\Delta x \Delta \nu$, or $\Delta y \Delta \nu$) that it passes through. This incidence-matrix approach automatically handles no-flux boundary conditions, and allows differential operators and PDE components to be constructed from products of incidence matrices and sparse diagonal matrices of metric information, as summarised in Table S6.

Table S6: **Matrices used in numerical simulations** (S50)

| Matrix | Size | Description |
|---|---|---|
| $\mathsf{A}$ | $J \times K$ | Incidence matrix mapping edges to vertices with directionality |
| $\mathsf{W}$ | $K \times K$ | Diagonal matrix of volumes around vertices (shaded grey area in Fig. S1), with factor of 1/2, 1/4, or 1/8 adjustments at domain boundary |
| $\mathsf{H}^v$ | $K \times K$ | Diagonal matrix of cisternal thickness over vertices |
| $\mathsf{H}^e$ | $J \times J$ | Diagonal matrix of cisternal thickness over edges ($= 0.5\overline{\mathsf{A}}\mathsf{H}^{\mathsf{v}}$) |
| $\mathsf{P}^\nu$ | $J \times J$ | Diagonal matrix to select only $\nu$ directed edges, with $P^\nu_{jj} = 1$ only if $j$ is a $\nu$-directed edge |
| $\mathsf{P}^x$ | $J \times J$ | Diagonal matrix to select only physical-space directed edges, with $P^x_{jj} = 1$ only if $j$ is not a $\nu$-directed edge |
| $\mathsf{L}$ | $J \times J$ | Diagonal matrix of edge lengths |
| $\mathsf{E}$ | $K \times K$ | Diagonal matrix of local enzyme concentration |
| $\mathsf{Q}^\perp$ | $J \times J$ | Diagonal matrix of perpendicular areas through which each edge passes, adjusted by appropriate factor at domain periphery |
| $\mathsf{R}$ | $K \times K$ | Diagonal matrix holding $R_{kk} = 1/(1 + \alpha_C H^v_{kk})$ |

By considering the divergence at vertices as the sum of fluxes along edges, we express a discretized form of (S39) as a coupled set of linear ordinary differential equations

$$\frac{\mathrm{d}\mathsf{C}}{\mathrm{d}\tilde{t}} = \mathsf{MC}, \quad \mathsf{M} \equiv \mathsf{RW}^{-1}\mathsf{A}^\top\mathsf{Q}^\perp \left(K_2K_4\mathsf{EP}^\nu\mathsf{L}^{-1}\mathsf{A} - \beta\mathsf{EP}^\nu\overline{\mathsf{A}}/2 + \mathcal{D}\mathsf{H}^e\mathsf{P}^x\mathsf{L}^{-1}\mathsf{A}\right). \tag{S50}$$

The matrices comprising the linear operator $\mathsf{M}$ are summarised in Table S6. $\mathsf{L}^{-1}\mathsf{A}$ is a generalised gradient operator, projected onto physical or polymerization space using $\mathsf{P}^x$ or $\mathsf{P}^\nu$. $\mathsf{W}^{-1}\mathsf{A}^\top\mathsf{Q}^\perp$ is a generalised divergence operator. The system is solved using an adaptive strong-stability-preserving Runge-Kutta scheme [1] from the Julia DifferentialEquations.jl library [2]. The time-varying matrix $\mathsf{E}$, capturing (S39b), must be recalculated at each timestep. We use an initial condition that is uniform in space, with a narrow Gaussian distribution in $\nu$, representing the early excess of unglycosylated cargo.

For the advective component we used central differencing, where the gradient operator is $\overline{\mathsf{A}}/2$ and $\overline{A}_{jk} \equiv |A_{jk}|$, rather than upstream differencing, where the gradient operator is $\overline{\mathsf{A}} - \mathsf{A}/2$, as this was found to produce a better-fit in comparison to analytic solutions, for the examples considered in the main text.

In cases of varying cisternal thickness, thickness profiles were derived from a Gaussian random field [3, 4] using the Julia `GaussianRandomFields.jl` package [5].

### S4 Uniform cisternal depth

Explicit solutions of the model can be derived for a cisterna that is spatially uniform, as we now demonstrate.

#### S4.1 Homogeneous enzyme, uniform depth

Suppose that $F_e(\mathbf{x}_\perp) = 1$ and $h(\mathbf{x}_\perp) = 1$. Then $\tilde{C}(\nu, \tilde{t})$ (dropping $\mathbf{x}_\perp$-dependence) satisfies, from (S39),

$$(1+\alpha_C)\tilde{C}_{\tilde{t}} + \tilde{E}\beta\tilde{C}_\nu = \tilde{E}K_2K_4\tilde{C}_{\nu\nu}, \quad \tilde{E}(\tilde{t}) = \left[1 + \frac{1}{K_2}\int_0^1 \tilde{C}(\nu,\tilde{t})\,\mathrm{d}\nu\right]^{-1}. \tag{S51a}$$

The conservation law (S37) ensures that $\int_0^1 \tilde{C}\,\mathrm{d}\nu = 1$ so that the level of enzyme is constant,

$$\tilde{E} = \frac{K_2}{1+K_2}. \tag{S52}$$

This reduces the problem to a linear advection-diffusion problem in $\nu$-space. A solution is illustrated in Fig. 2(a) of the main text. After a brief initial transient, and before the $\tilde{C}$-distribution encounters the boundary at $\nu = 1$ (i.e. for $\tilde{E}\beta\tilde{t} \ll 1+\alpha_C$), the solution is Gaussian (for the chosen parameters), and is well approximated by

$$\tilde{C}(\nu,\tilde{t}) \approx \sqrt{\frac{1+\alpha_C}{4\pi\tilde{E}K_2K_4(\tilde{t}-\tilde{t}_0)}}\exp\left[\frac{-\left\{(\nu-\nu_0)(1+\alpha_C) - \tilde{E}\beta(\tilde{t}-\tilde{t}_0)\right\}^2}{4\tilde{E}K_2K_4(1+\alpha_C)(\tilde{t}-\tilde{t}_0)}\right]. \tag{S53}$$

Here $\nu_0$ and $\tilde{t}_0$ are constants determined by the precise form of the initial condition; we evaluate them by fitting to simulation data (Fig. 2a), finding that both are small in magnitude. The mass of functional polymer rises as the Gaussian crosses the threshold $\nu = \phi$ (Fig. 2b), and is well approximated by

$$\tilde{M}_\phi(\tilde{T}_r) \equiv \int_\phi^1 \tilde{M}\,\mathrm{d}\nu = \pi\int_\phi^1 \tilde{C}(\nu,\tilde{T}_r)\,\mathrm{d}\nu = \frac{\pi}{2}\left[1 - \mathrm{erf}\left\{\frac{(\phi-\nu_0)(1+\alpha_C) - \tilde{E}\beta(\tilde{T}_r-\tilde{t}_0)}{\sqrt{4\tilde{E}K_2K_4(1+\alpha_C)(\tilde{T}_r-\tilde{t}_0)}}\right\}\right], \tag{S54}$$

so that

$$\mathcal{M}_\phi^* = \frac{\mathcal{C}\alpha_C}{2(1+\alpha_C)}\left[1 - \mathrm{erf}\left\{\frac{(\phi-\nu_0)(1+\alpha_C) - \tilde{E}\beta(\tilde{T}_r-\tilde{t}_0)}{\sqrt{4\tilde{E}K_2K_4(1+\alpha_C)(\tilde{T}_r-\tilde{t}_0)}}\right\}\right]. \tag{S55}$$

At large times, $\tilde{M}_\phi \to \pi$, as in Fig. 2(b) of the main text. We use the condition $\tilde{M}_\phi = \pi/2$ to identify $\tilde{T}_{r,50}$ from (S54) via

$$\tilde{E}\beta(\tilde{T}_{r,50} - \tilde{t}_0) = (\phi-\nu_0)(1+\alpha_C). \tag{S56}$$

Writing $u \equiv h_0/h_C$, (S56) can be rexpressed as

$$\frac{1}{\tilde{T}_{r,50}-\tilde{t}_0} = \frac{k_3\mathcal{S}}{k_1\mathcal{C}}\frac{\gamma N}{\phi-\nu_0}\frac{(1+u)(1-\Delta(1+\lambda u))}{[1+\gamma(1+u)](1+\lambda u)}. \tag{S57}$$

To understand the parameter dependence of production rate, we set $\tilde{t}_0 = 0$ and $\nu_0 = 0$ in (S57) for simplicity; the benefit of recovering a tractable prediction for the production rate is to offset the resulting quantitative error, which is modest in comparison to wider uncertainties in parameter values. Thus (S48) and (S57) become

$$\mathcal{P}^*_{50} \approx \frac{k_1 \mathcal{C}\mathcal{E}}{4|\Omega_\perp|N\phi} F(h_0/h_C) \text{ where } F(u) \equiv F(u;\lambda,\gamma,\zeta,\Delta) \equiv \frac{u\,[1-\Delta(1+\lambda u)]}{(1+u)(1+\zeta(1+\lambda u))(1+u+(1/\gamma))}. \tag{S58}$$

The function $F$ captures the dependence of cisternal depth on production rate. $F=0$ at $u=0$ and at the threshold identified by (S36). The location $u_{\max}$ at which $F=F_{\max}$ between these bounds is regulated by the parameters $\zeta$, $\lambda$, $\gamma$ and $\Delta$ (Table S5).

To illustrate one scenario, suppose that $\lambda \ll 1$ (with other parameters remaining order unity), implying that polymer adsorbs to the membrane much more weakly than the substrate ($h_C \ll h_S$). Then we can approximate $F$ as

$$F \approx \begin{cases} \dfrac{u(1-\Delta)}{(1+u)(1+\zeta)(1+u+(1/\gamma))} & \text{for} \quad u \sim O(1), \\ \dfrac{1-\Delta(1+\lambda u)}{u(1+\zeta(1+\lambda u))} & \text{for} \quad u \sim O(1/\lambda), \end{cases} \tag{S59}$$

with $F \approx (1-\Delta)/[(1+\zeta)u]$ in the overlap region for $1 \ll u \ll 1/\lambda$. Thus $F$ rises from zero to a maximum in the region for which $u \sim O(1)$, before decaying to zero in the region for which $u \sim O(1/\lambda)$. More specifically,

$$F_{\max} = \frac{1-\Delta}{(1+\zeta)\left[1+\sqrt{1+(1/\gamma)}\right]^2} \quad \text{for} \quad u = \sqrt{1+\frac{1}{\gamma}} \quad (\lambda \ll 1). \tag{S60}$$

In the original variables, the optimal cisternal thickness in this limit is

$$h_0 \approx h_C \sqrt{1+\frac{k_1\mathcal{C}}{2k_2|\Omega_\perp|}}, \tag{S61}$$

and the maximum production rate corresponding to (S60) is

$$\mathcal{P}^*_{50,\max} \approx \frac{\mathcal{C}\mathcal{E}}{4|\Omega_\perp|N\phi} \frac{k_1k_3\mathcal{S} - 2k_2k_4|\Omega_\perp|}{k_3\mathcal{S}+2k_2|\Omega_\perp|} \left[1+\sqrt{1+\frac{k_1\mathcal{C}}{2k_2|\Omega_\perp|}}\,\right]^{-2}. \tag{S62}$$

If, in addition, $\gamma \ll 1$, so that the polymer is abundant, then

$$\mathcal{P}^*_{50,\max} \approx \frac{\mathcal{E}k_2}{2N\phi} \frac{k_3\mathcal{S} - 2k_2k_4|\Omega_\perp|/k_1}{k_3\mathcal{S}+k_2|\Omega_\perp|} \qquad \text{at} \qquad h_0 \approx h_C \frac{k_1\mathcal{C}}{2k_2|\Omega_\perp|}. \tag{S63}$$

In this limit, the amount of available polymer $\mathcal{C}$ regulates the cisternal thickness at which the production rate is maximized, but not the maximum production rate itself.

An alternative limit is $\lambda \gg 1$ (implying that the substrate binds to the membrane much more weakly than the polymer, $h_S \ll h_C$) and $\zeta \gg 1$ (substrate is abundant). Then $F$ can be approximated by

$$F \approx \frac{u[1-\Delta(1+\lambda u)]}{\zeta(1+(1/\gamma))(1+\lambda u)} \quad \text{for} \quad u \sim O(1/\lambda). \tag{S64}$$

This function has a maximum satisfying

$$F_{\max} = \frac{1}{\lambda\zeta(1+(1/\gamma))}\left[1-\Delta^{1/2}\right]^2 \quad \text{at} \quad u = (1/\lambda)[\Delta^{-1/2}-1], \tag{S65}$$

or, in dimensional terms,

$$\mathcal{P}^*_{50} = \frac{k_1\mathcal{C}\mathcal{E}}{4|\Omega_\perp|N\phi}\frac{k_1\mathcal{C}}{k_1\mathcal{C}+2k_2|\Omega_\perp|}\frac{h_\mathcal{S}}{h_\mathcal{C}}\left[\left(\frac{k_3\mathcal{S}}{2k_2|\Omega_\perp|}\right)^{\frac{1}{2}} - \left(\frac{k_4}{k_1}\right)^{\frac{1}{2}}\right]^2 \text{ at } h_0 = h_\mathcal{S}\left[\left(\frac{k_1k_3\mathcal{S}}{2k_2k_4|\Omega_\perp|}\right)^{\frac{1}{2}} - 1\right]. \tag{S66}$$

In this limit, both the maximum production rate and the thickness at which this maximum is achieved are regulated by the amount of substrate.

#### S4.2 Variance

At the instant defined by (S56), $\tilde{C}$ in (S53) satisfies

$$\tilde{C}(\nu,\tilde{t}) = \sqrt{\frac{\beta}{4\pi K_2K_4(\phi-\nu_0)}}\exp\left[\frac{-\beta(\nu-\phi)^2}{4K_2K_4(\phi-\nu_0)}\right], \tag{S67}$$

a Gaussian distribution with variance (with respect to $\nu$) satisfying

$$\varsigma_{50}^2 = \frac{2K_2K_4(\phi-\nu_0)}{\beta}. \tag{S68}$$

The depth dependence of the variance is revealed by re-writing (S68) as

$$\varsigma_{50}^2 = \frac{2(\phi-\nu_0)}{N}\left(\frac{1}{\Delta(1+h_0/h_S)}-1\right)^{-1} \equiv \frac{2(\phi-\nu_0)}{N}\frac{\Delta(1+\lambda u)}{1-\Delta(1+\lambda u)}. \tag{S69}$$

Recalling the definition of $\Delta$ (Table S5), we find that the variance is regulated by the abundance, and adsorption properties, of substrate alone. It is smallest as $h_0 \to 0$, representing the intrinsic spread of the $\tilde{C}$ distribution arising because of multiple state transitions that are possible when starting with a large initial population of monomers. Eq. (S69) shows that the variance increases monotonically with cisternal depth, diverging as $h_0$ approaches the cut-off thickness at which the reaction fails to proceed. The variance with respect to polymer number can be written

$$(\varsigma_{50}^*)^2 = 2N(\phi-\nu_0)G(u), \quad G(u) \equiv \frac{\Delta(1+\lambda u)}{1-\Delta(1+\lambda u)}. \tag{S70}$$

This expression can be used to assess depth-dependence when the cisterna has uniform thickness; it is not suitable to be used when the cisterna has non-uniform depth, as $\tilde{C}$, when integrated over the domain, can be expected to become non-Gaussian, and the present approximation is not appropriate.

Table S7: **Representative parameter values.** L, T, M are arbitrary units of length, time and molecular number. Small or large parameters can be used to assess the effectiveness of the conditions in the right-hand column that are exploited to simplify the model. $h_{\text{cut-off}}$ is given by (S36). dim.=dimensional, nondim.=nondimensional.

| Input, dim. | Derived, dim. | Derived, nondim. | Condition |
|---|---|---|---|
| $\lvert\Omega_\perp\rvert = 10^4\mathrm{L}^2$ | $L_0 = 56.4\mathrm{L}$ | | |
| $\lvert\Omega\rvert = 10^4\mathrm{L}^3$ | $h_0 = 1\mathrm{L}$ | $h_0/L_0 = 1.8 \times 10^{-2}$ | Thin cisterna |
| $N = 100$ | | | Strong polymerization |
| $k_{Sa} = 1\mathrm{L/T}$ | | $D_S/(h_0 k_{Sa}) = 10^{-2}$ | Strong exchange |
| $k_{Sd} = 1\mathrm{T}^{-1}$ | $h_S = 2\mathrm{L}$ | $\alpha_S = 0.5$ | |
| $k_{Ca} = 0.1\mathrm{L/T}$ | | $D_C/(h_0 k_{Ca}) = 0.1$ | Strong exchange |
| $k_{Cd} = 1\mathrm{T}^{-1}$ | $h_C = 0.2\mathrm{L}$ | $\alpha_C = 5$, $\lambda = 0.1$ | Weak cargo affinity |
| $k_1 = 1\mathrm{L}^2/(\mathrm{MT})$ | | | |
| $k_2 = 2.5 \times 10^{-2}\mathrm{T}^{-1}$ | | $K_2 = 0.3$ | |
| $k_3 = 2.5 \times 10^{-2}\mathrm{L}^2/(\mathrm{MT})$ | | $K_3 = 2.5 \times 10^{-2}$ | |
| $k_4 = 1\mathrm{L}^2/(\mathrm{MT})$ | | $K_4 = 1$ | Limited dissociation |
| $\mathcal{C} = 10^4\mathrm{M}$ | $C_0 = 8.3 \times 10^{-2}\mathrm{M/L}^2$ | $\gamma = 5 \times 10^{-2}$ | Abundant monomer |
| | $C_b = 1\mathrm{M/L}^3$ | | |
| $\mathcal{S} = 10^5\mathrm{M}$ | $S_0 = 3.33\mathrm{M/L}^2$ | $\sigma = 40$, $\sigma K_3 = 1$ | Abundant substrate |
| | | $\zeta = 0.2$, $\Delta = 0.2$ | |
| | $S_b = 10\mathrm{M/L}^3$ | $\frac{S_b k_1 k_3 k_{Sa}}{k_2 k_4 k_{Sd}} = 10$ | Sufficient substrate |
| | $h_{\text{cut-off}} = 8\mathrm{L}$ | $\beta = 70$ | Forward reaction |
| $\mathcal{E} = 1.0\mathrm{M}$ | $E_0 = 5 \times 10^{-5}\mathrm{M/L}^2$ | $\varepsilon = 6 \times 10^{-4}$ | Scarce enzyme |
| | $T_0^* = 2.6 \times 10^8\mathrm{T}$ | | Production timescale |
| $D_C = 10^{-3}\mathrm{L}^2/\mathrm{T}$ | $(D_C\tilde{T}_0)^{1/2} = 0.124\mathrm{L}$ | $\delta_C = \delta_S = 3.1 \times 10^{-3}$ | |
| $D_S = 10^{-3}\mathrm{L}^2/\mathrm{T}$ | | $h_0/(D_C\tilde{T}_0)^{1/2} = 0.002$ | Rapid vertical diffusion |
| | | $\mathcal{D} = 204.2$ | |
| $\phi = 0.5$ | | | Functional threshold |

### S5 Parameter values and model assumptions

We choose L = 100 nm as a reference length (comparable to a cisternal thickness, see Fig. 1B of the main text), M = $10^3$ as a reference molecular number (representing a low number of membrane-bound enzymes in a cisterna), and T = $10^{-5}$ s as a reference time. With these reference quantities (which may be varied to apply in different contexts), we choose a set of parameters in column 1 of Table S7 (diffusion coefficients are comparable to 1 $\mu\mathrm{m}^2/s$) from which we derive parameters given in columns 2 and 3. These are chosen to approximately satisfy the model assumptions, listed in column 4.

There is considerable uncertainty in the biochemical parameters. They are presented to illustrate how the model might be used once more reliable parameter values are available. The assumptions are now discussed in turn.

- The cisternal aspect ratio $h_0/L_0$ is assumed small ($|\Omega|^2 \ll |\Omega_\perp^3|$, or $h_0/L_0 \ll 1$), to allow cross-cisternal averaging of bulk concentration fields.
- Formation of long polymers ($N \gg 1$) allows the discrete model to be approximated using a

continuous model. We assume $N = 100$.

- Adsorption of the cargo and substrate to the membrane is assumed to take place very rapidly in comparison to bulk diffusion (a strong exchange limit), ensuring that bulk and surface concentrations are always close to equilibrium.
- Adsorption strength can be assessed by the lengthscales $h_S$ and $h_C$. These can play a central role in determining the dependence of production rate on cisternal thicknesses. The three lengthscales are assumed to be of comparable magnitude ($h_S \sim h_C \sim h_0$) in Table S7. Weak affinity of the cargo for the membrane is represented by $\alpha_C \gtrsim 1$, implicating cisternal morphology in polymerization.
- The forward reactions are assumed to progress faster than the reverse reactions, enabling polymerization to proceed. This is facilitated by ensuring substrate is sufficiently abundant for its concentration to remain approximately constant ($\sigma \gg 1$), and for the cut-off thickness to exceed the cisternal thickness ($\beta > 0$, $h_0 < h_{\text{cut-off}}$).
- A further requirement is that complex formation and complex dissociation occur on comparable timescales ($\sigma K_3 \sim 1 \sim K_4$ with $\sigma \gg 1$)
- Enzyme is assumed to be sparse ($\varepsilon \ll 1$), making its abundance rate-limiting.
- The timescale $T_0^* = 2|\Omega_\perp| N^2 (K_2 + \sigma K_3)/(k_1 \mathcal{E})$ is the dimensional time corresponding to $\tilde{t} = 1$.
- The polymerisation reaction is assumed to be slower than the time for polymer and substrate molecules to diffuse across the thickness of the cisterna. The relevant reaction timescale is $T_0^*$, making the relevant condition $h_0 \ll (D_C T_0^*)^{1/2}$. This is comfortably satisfied by the chosen parameters.

#### S6 Two-chained polymer

The model can readily be extended to a polymer with two chains by replacing $C_n$ with $C_{n,m}$ and $Q_n$ with $Q_{n,m}$ in (S1) and including an additional reaction $Q^*_{n,m} + S^*_s \underset{k_4}{\overset{k_3}{\rightleftharpoons}} C^*_{n,m+1,s} + E^*$, allowing both chains of the polymer to grow at the same rate. This impacts terms involving $K_3$ and $K_4$ in the evolution equations (S24), which become $K_3 \sigma S (Q_{n-1,m} + Q_{n,m-1}) - 2K_4 C_{n,m} E (1 - \delta_{n1})(1 - \delta_{m1})$ in (S24a), for $n = 1, \ldots, N$, $m = 1, \ldots, N$; $2K_3 \sigma Q_{n,m} S - 2K_4 C_{n,m} E (1 - \delta_{n1})(1 - \delta_{m1})$ in (S24b); $-2K_3 \sigma Q_{n,m} S + K_4 E (C_{n+1,m} + C_{n,m+1})$ in (S24c) for $n = 1, \ldots, N-1$ and $m = 1, \ldots N-1$; and $-2K_3 Q_{n,m} S + (K_4 E/\sigma)(C_{n+1,m} + C_{n,m+1})$ in (S24d). The approximation $\sigma \gg 1$ leads to $S = 1$ to leading order. The approximation $\epsilon \ll 1$ leads to the equilibrium (S28) being modified to

$$(K_2 + 2\sigma K_3) Q_{n,m} = [C_n + K_4 (C_{n+1,m} + C_{n,m+1})] E. \tag{S71}$$

This allows appearances of $Q_{n,m}$ in the evolution equation for $C_{n,m}$ to be replaced with concentrations. Then, in the limit $N \gg 1$, the concentration distribution can be approximated by the continuous function $N^2 C_{n,m} = \tilde{C}(\nu, \mu, \mathbf{x}_\perp, \tilde{t})$ where $\mu = m/N$. Taylor-expanding, the analogue of

(S33) becomes

$$(1+\alpha_C h)\tilde{C}_{\tilde{t}}\frac{\mathrm{d}\tilde{t}}{\mathrm{d}t} = \frac{E}{N}\left(\tilde{C}_\nu + \tilde{C}_\mu\right)\frac{K_2K_4 - \sigma K_3}{K_2+\sigma K_3} + \frac{E}{2N^2}\frac{K_2K_4+\sigma K_3}{K_2+\sigma K_3}\left(\tilde{C}_{\nu\nu}+\tilde{C}_{\mu\mu}\right) + \frac{E\sigma K_3K_4}{N^2(K_2+2\sigma K_3)}\left(\tilde{C}_{\nu\nu}+\tilde{C}_{\mu\mu}-2\tilde{C}_{\mu\nu}\right) + \alpha_C\delta_C\nabla_\perp\cdot(h\nabla_\perp\tilde{C}). \tag{S72}$$

The term $\tilde{C}_{\nu\nu}+\tilde{C}_{\mu\mu}$ is an isotropic diffusion in $(\nu,\mu)$-space, whereas the following term involves $(\partial_\nu-\partial_\mu)^2\tilde{C}$, representing diffusion along $\mu+\nu$ =constant, which suppresses variations of $\tilde{C}$ having fixed total molecular weight. This motivates a simplified solution of the form $\tilde{C}(\mathbf{x}_\perp,\nu,\mu,\tilde{t}) = \breve{C}(\mathbf{x}_\perp,\rho,\tilde{t})$, where $\rho=\mu+\nu$, satisfying

$$(1+\alpha_C h)\breve{C}_{\tilde{t}}\frac{\mathrm{d}\tilde{t}}{\mathrm{d}t} = \frac{E}{N}\frac{K_2K_4-\sigma K_3}{K_2+\sigma K_3}2\breve{C}_\rho + \frac{E}{2N^2}\frac{K_2K_4+\sigma K_3}{K_2+\sigma K_3}2\tilde{C}_{\rho\rho} + \alpha_C\delta_C\nabla_\perp\cdot(h\nabla_\perp\tilde{C}). \tag{S73}$$

Relative to the single chain model, this corresponds to a doubling of the advective and diffusive terms in polymerization space.
